# Convergent deployment of ancestral programs during the evolution of mammalian flight membranes

**DOI:** 10.1101/2022.12.06.518272

**Authors:** Charles Y. Feigin, Jorge A. Moreno, Raul Ramos, Sarah A. Mereby, Ares Alivisatos, Wei Wang, Renée van Amerongen, Jasmin Camacho, John J. Rasweiler, Richard R. Behringer, Bruce Ostrow, Maksim V. Plikus, Ricardo Mallarino

## Abstract

Lateral flight membranes, or patagia, have evolved repeatedly in diverse mammalian lineages. While little is known about patagium development, its recurrent evolution may suggest a shared molecular basis. By combining transcriptomics, developmental experiments, and mouse transgenics, we demonstrate that lateral WNT5A expression in the marsupial sugar glider (*Petaurus breviceps*) promotes the differentiation of its patagium primordium. We further show that this function of WNT5A reprises ancestral roles in skin morphogenesis predating mammalian flight and has been convergently employed during patagium evolution in eutherian bats. Moreover, we find that many genes involved in limb development have been re-deployed during patagium outgrowth in both the sugar glider and bat. Taken together, our findings reveal that deeply conserved molecular toolkits underpin the evolutionary transition to flight in mammals.

## Introduction

Flight has arisen as many as seven times among extant mammals. This includes bats which evolved powered flight, as well as six instances of unpowered flight, or gliding, in diverse taxa such as colugos, flying squirrels and marsupial possums (*1-4*). Each flying lineage has evolved at minimum a lateral flight membrane, or patagium between the fore- and hindlimbs. This structure acts as an airfoil, generating lift and permitting directional movement during flight (**Fig. 1A**) (*5*). Notably, these independent derivations of a lateral patagium have originated at a wide range of evolutionary divergences, from ∼30 million years among some possums to the massive gulf of ∼160 million years that separate marsupial and eutherian mammals (*2, 6, 7*). Presently, little is known about the regulation of lateral patagium development in these species. However, the repeated evolution of this structure among diverse mammals suggests that it may be derived from shared ancestral developmental programs, whose barriers to evolutionary redeployment in the skin may be small and easily overcome. Therefore, resolving the molecular basis of patagium formation may yield generalizable findings about mammalian development and insights into how phenotypic novelty emerges from evolutionarily conserved programs.

**Fig. 1.**
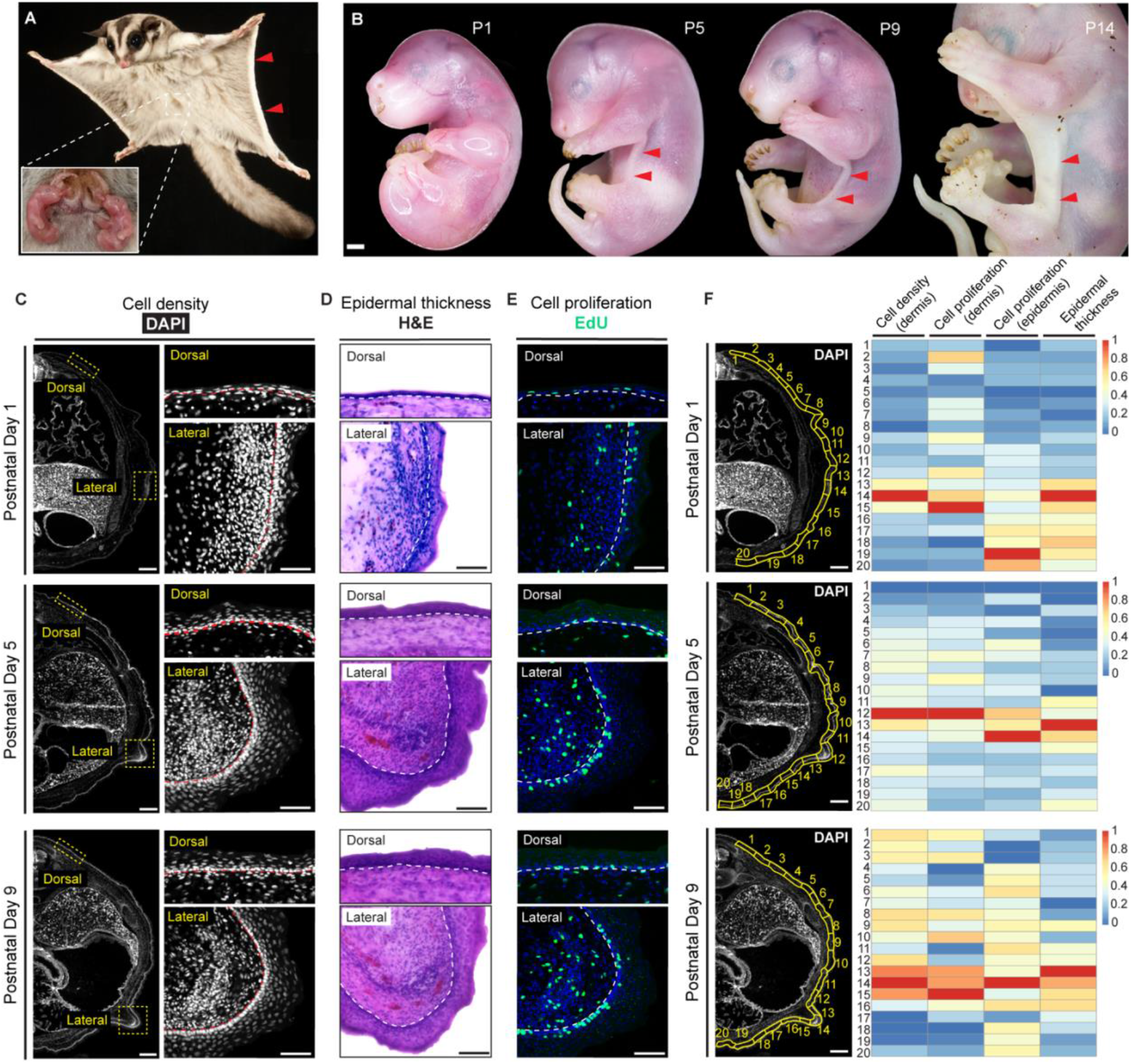
Development of the lateral patagium in the sugar glider. (A) An adult sugar glider in flight. Red arrows indicate the lateral patagium. Inset image shows pouch young laying atop the everted maternal pouch. (B) Development of sugar glider pouch young during the first two weeks postpartum. Red arrowheads indicate position of the lateral patagium. (C) DAPI stained transverse sections illustrating elevated cell density in the patagium primordium relative to dorsal skin. (D) Hematoxylin and Eosin (H&E) stained transverse sections showing elevated epidermal thickness in the patagium primordium relative to dorsal skin. (E) Transverse sections showing elevated cell proliferation in the patagium primordium relative to dorsal skin, as indicated by higher EdU incorporation. (F) Quantification of the histological properties of sugar glider skin along the dorsal-ventral axis in transverse sections. Cell density of the dermis, thickness of the epidermis, and cell proliferation of both tissue layers were quantified within the illustrated regions of interest (ROIs), outlined in yellow and presented as a heatmap. Dotted lines in (C-E) delineate the dermis-epidermis boundary. Scale bars: 1mm in (B); 500μm (zoomed out) and 100μm (zoomed in) in (C); 100μm in (D); 100μm in (E); and 500μm in (F). Photo credit in (A): Joe McDonald.

To uncover the mechanisms governing lateral patagium formation, we have developed genomic and experimental resources in a novel model species, the marsupial sugar glider (*Petaurus breviceps*). Like other marsupials, sugar gliders give birth to extremely altricial young that are developmentally comparable to the fetuses of eutherian mammals (*8, 9*). These neonates complete much of their development *ex utero* inside of their mother’s pouch and are thus referred to as pouch young (**Fig. 1A**) (*8*). Notably, the patagium is not discernable externally at birth in sugar glider pouch young (**Fig. 1B**). Rather, outgrowth of this structure begins several days postpartum. Together, the experimental accessibility of the skin, the postnatal formation of the patagium, and the sugar glider’s amenability to captive husbandry make this a very tractable model system to uncover how spatially patterned morphological structures originate.

## Results

### Characterization of lateral patagium formation in the sugar glider

We first characterized lateral patagium development in the sugar glider during the first two weeks postpartum. We observed that pouch young first show external evidence of the patagium at approximately 4 days postnatally (P5), in the form of a lateral ridge, most visible at the axilla. This ridge then extends outward over several weeks, gradually filling the inter-limb space (**Fig 1b**). Next, we examined the development of the patagium histologically, comparing it to neighboring skin along the dorso-ventral axis in transverse sections (**Fig. 1, C to F**). At P1, prior to the initiation of outgrowth, we observed an uneven distribution of nuclei in the dermal mesenchyme (indicated by DAPI staining), with higher cell density in the inter-limb lateral skin that constitutes the patagium primordium, relative to more dorsal and ventral skin regions (**Fig. 1, C and F**). This disparity was subsequently maintained during patagium outgrowth. Of particular note, we observed a central condensate of dermal mesenchyme marking the position of the nascent patagium primordium (**Fig. 1C**). This condensate was just discernable at P1 but became more prominent before the initiation of patagium outgrowth. Interestingly, we also observed substantial thickening (hyperplasia) in the epidermis overlying this mesenchymal condensate, a phenotype which declined with greater dorsal and ventral distance from the patagium primordium (**Fig. 1, D and F**). Intraperitoneal injections of pouch young with the proliferation marker 5-ethynyl-2’-deoxyuridine (EdU) revealed elevated cell proliferation in the dermal layer of the patagium primordium relative to neighboring skin (**Fig. 1, E and F; movie S1**). This increase was especially pronounced in the mesenchymal condensate, as well as in the epidermal basal layer immediately above it as development progressed, suggesting that proliferation may contribute to the establishment of observed morphological phenotypes in the dermal and epidermal layers. Together, these observations indicate that patagium formation is comprised of an initial phase of histological differentiation of the primordium, marked by increased cell proliferation, dermal condensation and epidermal hyperplasia in the patagium primordium, followed by a subsequent outgrowth phase which ultimately produces the flight membrane. Given this, we next sought to investigate the molecular basis of these developmental processes.

### The sugar glider genome

To facilitate functional genomic interrogations of patagium formation, we generated a draft genome assembly for the sugar glider using linked-read technology (10X Genomics) (*10*). The resulting assembly was ∼3.4 gigabases in length, had a scaffold N50 of ∼28 megabases, and was highly contiguous, with less than 2% gaps (**table S1**). Recovery of complete, single-copy mammalian BUSCOs (Benchmarking Universals Single-Copy Orthologs) was 86.5%, comparable to that of other recently released genomes of related diprotodont marsupials (**fig. S1 and table S2)**. We next annotated sugar glider genes by integrating multiple lines of transcript and protein alignment evidence, as well as *ab initio* gene predictions (*11-13*). This yielded 26,489 gene models, the identities of which were inferred by conditional reciprocal best blast searches against koala (*Phascolarctos cinereus*) RefSeq transcripts (*12, 14*). Our sugar glider draft assembly represents a valuable new resource for functional and evolutionary genomic studies.

### Localized elevation of Wnt-signaling in the patagium primordium

Given that patagium outgrowth is anticipated by localized histological changes in the lateral skin, we next sought to identify potential molecular drivers of this differentiation. To accomplish this, we performed RNA sequencing (RNA-Seq) on patagium skin samples from sugar glider pouch young spanning the first three weeks of postnatal development (N = 34; **table S3**) and used weighted gene correlation network analysis (WGCNA) to predict modules of genes with highly correlated expression levels across samples (*15*). Such correlation modules can reflect functional interactions between their constituent genes, or roles in concurrent biological processes. WGCNA analysis predicted 13 correlation network modules, each showing distinct patterns of module eigengene expression (a summary metric for the relative expression levels of its constituent genes) over the course of patagium development (**Fig. 2A**). Of particular interest was patagium module 8 (hereafter, PM8), a relatively small module containing only ∼4.32% of patagium-expressed genes **(Fig. 2A and table S4**). PM8 exhibited a stronger inverse correlation (r = -0.77, p < 0.001) between its eigengene expression and animal weight (a proxy for age) than did other modules. Thus, the period of highest expression for its constituent genes corresponded closely with the onset of histological differentiation in the patagium primordium. Notably, Gene Ontology analysis indicated that PM8 was highly enriched for Wnt-signaling genes (**tables S5 and S6**) (*16*). These data indicated that genes with roles in Wnt-signaling belonging to PM8 may help to drive morphological differentiation of the patagium primordium prior to its later outgrowth.

**Fig. 2.**
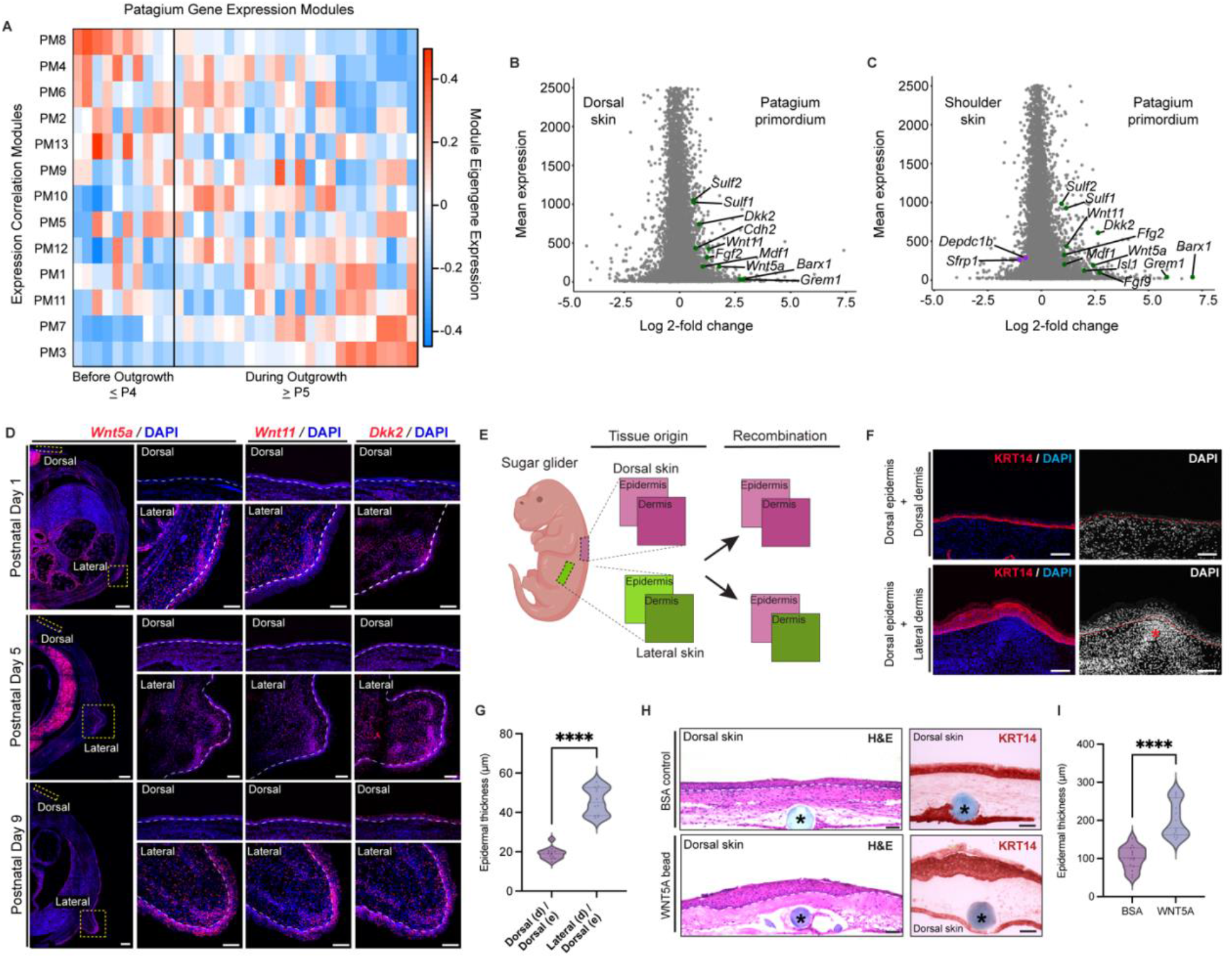
A distinct transcriptomic landscape drives the establishment of early patagium phenotypes. (A) Heatmap displaying eigengene expression of WGCNA modules across early patagium development. Columns represent individuals, sorted by increasing weight. Rows represent modules, sorted by their correlation with sample age. Cell color represents module eigengene expression. (B-C) Genes differentially expressed between the patagium primordium and dorsal (B) or shoulder skin (C). Labeled genes are those that were both differentially expressed in the patagium primordium relative to comparison skin regions (upregulated shown in green, downregulated shown in purple), and belonged to the enriched Wnt-signaling Gene Ontology term (GO:0030111). (D) *In situ* hybridizations of *Wnt5a, Wnt11* and *Dkk2*. (E) Schematic of tissue recombination experiments. (F) Explants stained for KRT4 and DAPI. Asterisk denotes the mesenchymal condensate. (G) Quantification of epidermal thickness in recombination experiments. (H) Hematoxylin-eosin (H&E) and KRT14 stained, bead-implanted explants. Asterisk shows implanted bead. (I) Quantification of epidermal thickness in bead-implanted explants. Statistical significance in panels G (**** P < 0.0001; N = 5) and I (**** P < 0.0001; N = 4) was assessed using a general mixed effects model ANOVA test. Dotted lines in (D) and (H) delineate the dermis-epidermis boundary. Scale bars: 400μm (zoomed out) and 100μm (zoomed in) in (D); 100μm in (F); and 200μm in (H).

If localized changes in Wnt-signaling contribute to the early differentiation of the patagium as our data suggest, we should expect this to be reflected by differences in gene expression levels relative to neighboring, non-patagium skin. Therefore, we compared the transcriptome of the skin constituting the patagium primordium to that of stage-matched samples of neighboring dorsal skin, prior to the initiation of patagium outgrowth (N = 6). Differential gene expression analysis showed that indeed, the expression levels of many of the Wnt-signaling genes belonging to PM8 were significantly greater in the patagium primordium relative to the dorsal skin (**Fig. 2B, table S7**) (*17*). Upregulated genes included those encoding the Wnt-family ligands *Wnt5a* and *Wnt11*, as well as the Wnt-signaling modulator *Dkk2*. These three secreted factors are generally associated with activation of non-canonical Wnt-signaling and/or suppression of canonical Wnt/β-catenin signaling (*18-20*). While this observation is consistent with a distinct Wnt-signaling environment during the differentiation of the patagium primordium, these results may alternatively be explained by more general characteristics of lateral skin, rather than of the patagium specifically. To explore this possibility, we sequenced RNA from stage-matched samples of shoulder skin (N = 6), which sits on the opposite side of the forelimb from the patagium and occupies the same coronal plane, thus constituting a non-patagium lateral skin region (**table S8**). As with our comparisons against dorsal skin, roughly the same complement of PM8 Wnt-signaling genes was found to be upregulated in the patagium primordium relative to shoulder skin. Indeed, several of these genes (e.g., *Wnt5a, Dkk2, Sulf1* and *Sulf2*) showed even larger fold increases (**Fig. 2C**). This analysis further supports localized elevation of Wnt-signaling being a distinct molecular feature of the inter-limb lateral skin comprising the patagium primordium.

Wnt-signaling primarily regulates morphogenesis through intercellular communication, mediated by diffusible factors (*21*). Given this, we next characterized the spatial distribution of cells expressing the most strongly upregulated PM8 Wnt-signaling factors, *Wnt5a, Wnt11* and *Dkk2*. Fluorescence *in situ* hybridization showed that expression of these genes was strong in the mesenchymal condensate of the patagium primordium and extended into the overlying basal layer of the thickened epidermis above (**Fig. 2D**). By contrast, few cells expressed these genes in the dorsal skin. The spatial and temporal expression pattern of Wnt-signaling genes further indicated that this pathway could play roles in driving differentiation of the patagium primordium. Therefore, we next sought to test for such roles through functional studies.

### WNT5A drives epidermal hyperplasia in the sugar glider’s patagium primordium

The Wnt-expressing mesenchymal condensate of the patagium primordium is overlaid by a region of epidermal hyperplasia (**Fig. 1, C to F and Fig. 2D**). This spatial correspondence led us to hypothesize that the mesenchymal condensate may act as a signaling center, controlling thickness of the overlying epidermis. To test this, we performed epidermal-dermal tissue recombinations between patagium and dorsal skin of sugar glider pouch young in an explant culture system. Briefly, we isolated samples of patagium and dorsal skin from pouch young just prior to the initiation of patagium outgrowth at P4, when the mesenchymal condensate was most prominent. We then enzymatically dissociated the two tissue layers and recombined segments of dorsal epidermis with patagium dermis (N = 5), or dorsal epidermis with dorsal dermis as a control (N = 5) and cultured them as explants for 72hrs (**Fig. 2E**). Immunohistochemistry (IHC) for keratin 14 (KRT14), a marker of the basal epidermal layer, showed that dorsal epidermises cultured on patagium dermises increased in thickness relative to controls (**Fig. 2, F and G**). This phenotype resembled differences in epidermal thickness observed between the patagium and dorsal skin *in vivo* (**Fig. 1, C and F**) and is consistent with our hypothesis that signals in the condensed mesenchyme of the patagium primordium control the thickness of overlying epidermis.

Having demonstrated that signals from condensed dermal mesenchyme do indeed drive epidermal hyperplasia, we next asked whether this effect may be regulated by Wnt-signals in particular. Among genes encoding Wnt-family ligands in PM8, *Wnt5a* was the mostly highly upregulated in the patagium primordium compared to neighboring skin regions (**Fig. 2, B and C**). The WNT5A ligand is a well-known morphogen that regulates the development of multiple structures during embryogenesis and plays a key role driving distal-specific gene expression (*22*). For instance, WNT5A expression in dermal fibroblasts induces expression of keratin 9 (*Krt9*), an epidermal differentiation gene expressed in acral skin regions such as the palms and soles (*23*). Moreover, dermal *WNT5A* upregulation has been implicated in psoriasis, a chronic skin disease characterized by pathologic epidermal hyperplasia (*24, 25*). We therefore tested the capacity of WNT5A to drive epidermal hyperplasia in sugar glider skin by soaking beads in recombinant WNT5A or bovine serum albumin (BSA) as a control and implanting them into the dermis of dorsal skin explants taken from pouch young. Consistent with our hypothesis, explants cultured with dermally implanted, WNT5A-soaked beads showed a marked increase in epidermal thickness compared to BSA-treated control explants (**Fig. 2, H and I**), resembling the phenotype produced by recombining dorsal epidermis with patagium dermis (**Fig. 2, F and G**). This finding supports a key role for locally elevated WNT5A in mediating dermal control of epidermal hyperplasia in the earliest stages of patagium differentiation.

### WNT5A promotes dermal condensation in mammalian skin

WNT5A is involved in the formation of diverse mesenchymal condensates during animal development, including chondrogenic condensates during bone development in the distal limb (*26*) and dermal condensates (precursors of the dermal papillae) during hair follicle formation (*27*). Therefore, we next asked whether WNT5A may also contribute to the formation of the mesenchymal condensate observed in the patagium primordium. To this end, we leveraged the mature genetic tools available in the laboratory mouse (*Mus musculus*), by generating transgenic lines in which *Wnt5a* could be inducibly upregulated (*R26rtTA;tetO-Wnt5a*) by treating animals with doxycycline (*28, 29*). Examination of mouse skin at postnatal day 37 (P37) revealed that induction of *Wnt5a* over-expression for 7 days drove a marked increase in dermal cell density in double-transgenic mice (*R26rtTA*^*HET*^*;tetO-Wnt5a*^*HET*^; N = 4) compared to doxycycline-treated control littermates (*R26rtTA*^*WT*^*;tetO-Wnt5a*^*HET*^; N = 4) (**Fig. 3, A and B**). Relative to *Wnt5a* expression in the patagium primordium of sugar glider pouch young, our transgenic mouse model drives relatively uniform upregulation of *Wnt5a* throughout the skin. Thus, no discrete, lateral mesenchymal condensate was formed. Nonetheless, this finding supports WNT5A’s ability to drive mesenchymal condensation in the skin of mammals, consistent with such a role in the patagium primordium. Interestingly, as occurred in our sugar glider explant experiments, we also observed epidermal hyperplasia in double-transgenic mice compared to controls, as evidenced by histology and IHC for KRT14, further supporting the role of WNT5A in driving this phenotype (**Fig. 3, C and D**). Taken together, our experiments indicate that localized Wnt-signaling, mediated in part by WNT5A, coordinates the differentiation of both dermal and epidermal layers in the patagium primordium of the sugar glider.

**Fig. 3.**
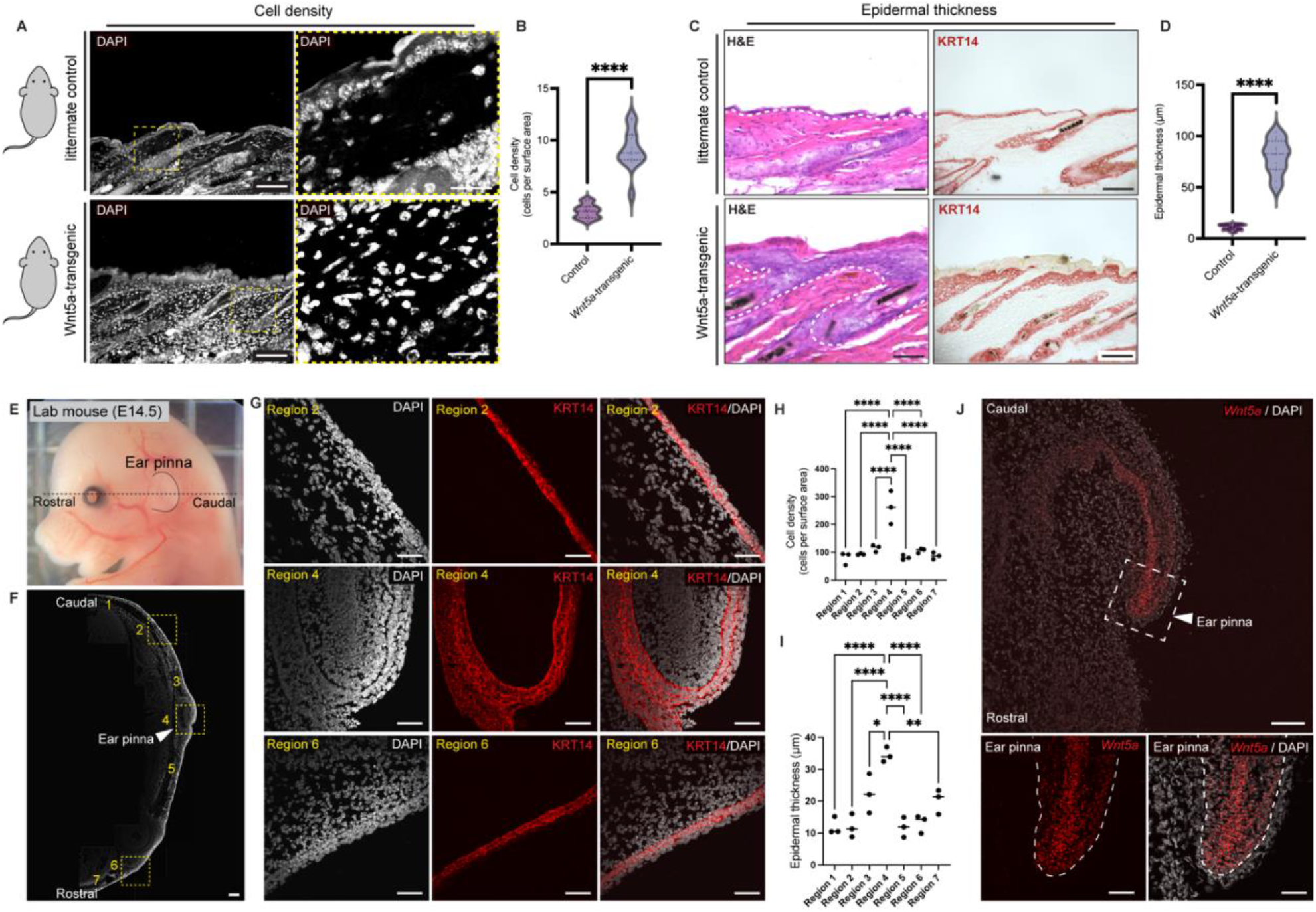
*Wnt5a*’s role in skin morphogenesis is ancestral in mammals. (A) DAPI-stained transverse sections showing dermal cell density in transgenic mice and controls. Yellow box denotes zoomed-in region. (B) Quantification of cell density in transgenic mice and controls. (C) Hematoxylin-Eosin (H&E) and KRT14 stained mouse skin from transgenic mice and controls. (D) Quantification of epidermal thickness in transgenic mice and controls. (E) E14.5 mouse, showing the early ear pinna. Dotted line shows the rostro-caudal axis along which sections were taken. (F) ROIs used for quantifying dermal cell density and epidermal thickness. (G) Sections stained with DAPI and KRT14. (H-I) Quantification of cell density (H) and epidermal thickness (I) in E14.5 mouse pinna. (J) *In situ* hybridization for *Wnt5a* in E14.5 mouse pinna. Statistical significance in panels (B and D) (N = 4; **** P < 0.001) and in panels (H and I) (**** P < 0.0001; *** P < 0.001; ** P < 0.01; *P < 0.05; N = 3) was assessed using a general mixed effects model ANOVA test. Post-hoc pairwise comparisons indicated that Region 4 was significantly different from other regions. All other comparisons were statistically not significant and are not displayed. Dotted lines in (C) and (J) delineate the dermis-epidermis boundary. Scale bars: 200μm (zoomed out) and 50μm (zoomed in) in (A); 100μm in (C); 200μm in (F); 50μm in (G); 100μm (zoomed out) and 50μm (zoomed in) in (J).

### Convergent deployment of an ancestral WNT5A function in skin morphogenesis

It is notable that induction of *Wnt5a* is able to drive phenotypes reminiscent of those observed during early patagium differentiation in the sugar glider in the skin of a mouse, a species which is fully terrestrial and last shared a common therian mammal ancestor with sugar gliders ∼160 million years ago (*7*). This observation may indicate an ancestral origin for WNT5A’s function in skin morphogenesis that long-precedes the evolution of mammalian flight. If this is the case, roles for WNT5A in condensation of dermal mesenchyme and epidermal thickening should be discernable in the skin of other mammalian structures unrelated to flight. We therefore examined the skin of a distinct structure common to therian mammals: the pinna of the ear. To this end we collected laboratory mouse samples at embryonic day 14.5 (E14.5; N = 3), early in the formation of the pinna and after the initial differentiation of the epidermis (**Fig. 3, E and F**) (*30*).

Quantification of cell density by DAPI showed that, like the patagium in the sugar glider, the mouse pinna had a marked increase in mesenchymal density relative to neighboring head skin along the rostro-caudal axis, corresponding to tissue that will form ear dermis and ear cartilage later in development (**Fig. 3, F to H**). IHC for KRT14 showed that additionally, epidermis overlying the distal pinna exhibited hyperplasia relative to neighboring skin of the head **(Fig. 3, F, G and I**). Given this, we next performed fluorescent *in situ* hybridization to examine the spatial distribution of *Wnt5a*-expressing cells. This revealed that *Wnt5a* was robustly expressed within the mesenchyme of the pinna, with strongest expression near the distal tip where epidermal thickness peaked (**Fig. 3J**). Moreover, we found that *Wnt5a* expression was nearly absent from neighboring head skin (**Fig. 3J**). These observations support an ancestral origin for WNT5A’s function in mammalian skin morphogenesis.

Given that the role of WNT5A in skin morphogenesis appears to predate the origin of flight, this role should, in principle, be part of the shared, ancestral developmental genetic toolkit of distantly related mammals. This raised the question of whether WNT5A might also be involved in the early stages of patagium formation in other flying mammals that have independently evolved a lateral patagium. To explore this possibility, we acquired field-collected embryonic samples of the microbat *Carollia perspicillata* within a narrow developmental window (Stages 15-17). This period corresponds to the early external formation of the lateral patagium, which in bats is referred to as the plagiopatagium to distinguish it from several other developmentally distinct membranes comprising the bat wing. We then performed histological analyses and compared their results to those in the sugar glider (**Fig. 4, A and B**) (*31*).

**Fig. 4.**
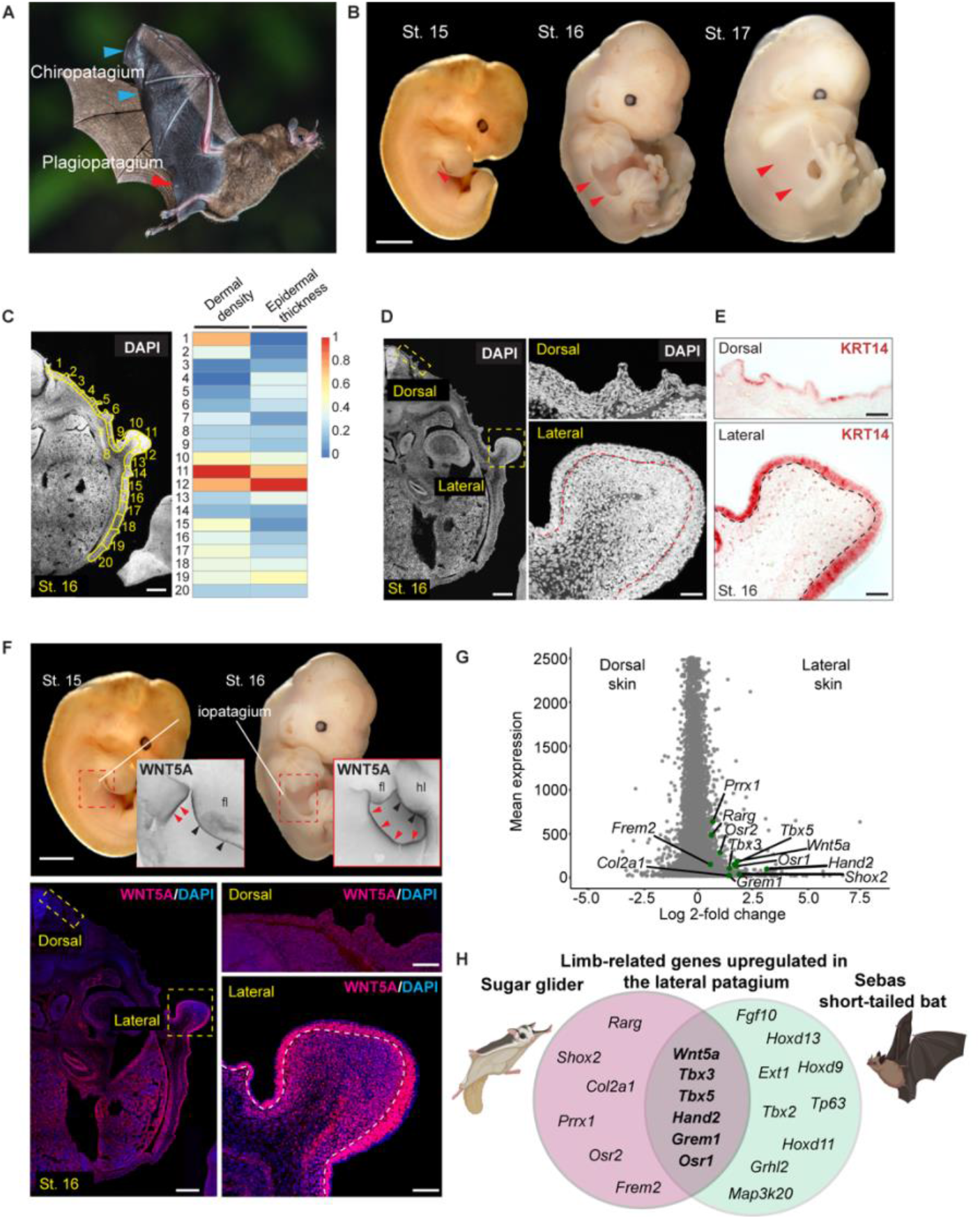
Evolutionary convergence between the lateral patagia of sugar gliders and bats. (A) The plagiopatagium (red arrowheads) of Seba’s short-tailed bat is shown in an adult animal and (B) in embryonic stages. This structure differs from other patagia in the bat wing, such as the chiropatagium (blue arrowheads). (C) Quantification of dermal cell density and epidermal thickness along the dorsal-ventral axis in an embryonic bat (St. 16) within multiple ROIs (highlighted in yellow) and illustrated in a heat map. (D and E) Bat plagiopatagium relative to dorsal skin in transverse sections, with cell density visualized by DAPI (D) and epidermal thickness visualized by IHC for KRT14 (E). (F) Whole-mount in situ hybridization of embryonic bats showing Wnt5a expression in the developing plagiopatagium (red arrowheads) relative to dorsal skin. Wnt5a is also detected in the limbs (black arrowheads) (fl: forelimbs; hl: hindlimbs). (G) Volcano plot of genes differentially expressed between the sugar glider patagium and the neighboring dorsal skin during its outgrowth. Patagium-upregulated genes belonging to limb development Gene Ontology terms are labeled in green. (H) Venn diagram showing genes with known roles in limb outgrowth and patterning convergently-upregulated in the developing lateral patagia of sugar gliders and bats. Photo credit in (A): Andrew Morffew.

Transverse sections from the developing bat plagiopatagium revealed striking similarities with the patagium of the sugar glider. Relative to neighboring skin along the dorso-ventral axis, the bat plagiopatagium showed a marked increase in mesenchymal cell density (**Fig. 4, C and D**). Moreover, we observed a corresponding thickening of the overlying epidermis (**Fig. 4, C and D**). These phenotypes strongly resembled those driven by WNT5A in sugar glider explants and in *Wnt5a*-overexpressing mice. Notably though, we observed that in contrast to the sugar glider, no central mesenchymal condensate was discernable. Rather, a broader area of mesenchyme within the nascent plagiopatagium was found to be compacted relative to neighboring skin. Interestingly, IHC revealed that this broader region of condensation corresponded with a similarly broad expression pattern of WNT5A within the developing plagiopatagium, and an extension of its expression into the overlying epidermis (**Fig. 4, E and F**). As indicated by our *Wnt5a*-overexpressing transgenic mice (**Fig. 3, A to D)**, a broader expression domain of WNT5A in the bat plagiopatagium may help explain why this structure exhibits a broader region of mesenchymal condensation (**Fig. 4F)** rather than a single, discrete mesenchymal condensate comparable to that seen in the sugar glider **(Fig. 2D)**.

Taken together, our findings demonstrate a key role for altered expression of Wnt-signaling genes, particularly *Wnt5a*, in driving early differentiation of the lateral patagium in the sugar glider. Moreover, we find evidence that this function is ancestral to mammals and has been convergently employed during the evolution of flight in the eutherian bats in spite of their large evolutionary separation from marsupials.

### Convergent deployment of limb development genes in the lateral patagia

Specification of the lateral patagium to the same coronal plane as the limbs is a functional necessity in both sugar gliders and bats, as a physical connection between these two morphological structures is ultimately required to regulate tension of the aerodynamic surface and to control direction during flight (*32*). In addition to a limb-like lateral position, it is notable that the lateral patagium’s outgrowth also proceeds in a manner reminiscent to that of the limbs in both species. The patagium first becomes visible externally as lateral ridge, which then extends outward (**Fig. 1B**). This ridge may represent a bud, akin to that of the early limbs, albeit a highly elongated one (*33*). This pattern of bud outgrowth contrasts markedly with many other skin membranes involved in tetrapod locomotion, such as the webbing found between the toes of water birds and turtles, as well as the chiropatagia of bat wings. Such interdigital membranes arise largely through localized suppression of apoptosis in the early limb bud, rather than as a distinct, novel outgrowth of skin (*34-36*).

The apparent developmental analogies between the lateral patagium and limbs in terms of their shared coronal plane and bud-like outgrowth suggest a potentially deeper molecular and evolutionary relationship between these structures. Indeed, WNT5A which contributes to the early differentiation of the patagium in both sugar gliders and bats is also a known driver of tetrapod limb formation (*22*). To explore this possibility, we next tested for upregulation of limb morphogenesis genes in the lateral patagia of sugar gliders and bats during their outgrowth phase, relative to stage-matches samples of neighboring dorsal skin (**tables S3, S9 and S10**). Consistent with a developmental-genetic relationship with the limbs, we observed significant enrichment of multiple limb development Gene Ontology terms among genes upregulated in the patagia of both species, including numerous transcription factors and morphogens (**Fig. 4G, fig. S2 and tables S9 to S14**). Most strikingly however, we observed a complement of genes upregulated in both species: *Wnt5a*, the BMP antagonist *Grem1*, and the transcription factors *Tbx5, Tbx3, Hand2* and *Osr1*; (**Fig. 4H and table S15**). *Tbx5* is necessary for the initiation of forelimb outgrowth while TBX3 helps to define limb bud width and participates in a transcriptional network with HAND2 to regulate anterior-posterior patterning (*37-39*). *Grem1* encodes a BMP/TGFβ inhibitor, which similarly participates in anterior-posterior digit patterning (*40*). *Osr1* by contrast identifies interstitial muscle connective tissues in the limb and is known to interact with *Tbx5* in other developmental contexts (*41, 42*).

Convergent activation of limb development genes in the sugar glider and bat reinforces the notion that spatial patterning and morphogenesis of the patagium may involve at least partial evolutionary redeployment of processes also involved in limb formation. Interestingly though, this transcriptional convergence is limited, with more uniquely redeployed limb development genes in each species than those that are shared between them (**Fig. 4H**). Nonetheless, given that the limbs are ancestral to all tetrapods, re-deployment of limb morphogenesis genes may present a parsimonious explanation for the recurrent evolution of the patagium across distantly related mammalian lineages.

## Discussion

Flight has evolved many times among animals. Across lineages, the core structures that permit this form of locomotion vary widely. In birds, wings evolved through modifications to the ancestral tetrapod forelimb, including changes in digit number and morphology as well as the derivation of specialized skin appendages (i.e., flight feathers). In insects, recent evidence has suggested that wings are derived from ancestral leg segments and are homologous to the tergal plate found in crustaceans (*43*). In these lineages, dedicated flight structures are thought to have evolved only once. By contrast, specialized flight structures appear to have arisen at least seven times independently among extant mammals, at a wide range of evolutionary divergences. Notably, a universal feature of both powered and unpowered flight in mammals is the growth of a novel, membranous lateral patagium connecting the limbs. The recurrent evolution of lateral patagia in diverse mammalian lineages suggests that it may reflect shared developmental programs that predate flight.

Here, we show that formation of the patagium primordium in the marsupial sugar glider is marked by upregulation of Wnt-signaling genes. We demonstrate that WNT5A in particular contributes to both condensation of dermal mesenchyme and thickening of the overlying epidermis in this region. Our studies in eutherian laboratory mice further show that this role for WNT5A is likely conserved in mammals, controlling skin morphogenesis in very distinct structures, such as the pinna of the ear. Furthermore, we find evidence that this ancestral function for WNT5A in skin development has been convergently employed during the evolution of the lateral plagiopatagium in bats, driving similar phenotypes in both dermal and epidermal skin layers in spite of their ∼160 million year separation (*7*).

While enhanced signaling through the WNT5A ligand appears to play a role in driving early phenotypes shared between the bat and sugar glider, it is less clear what developmental requirement there might be for epidermal thickening or dermal condensation during patagium formation. It is perhaps most plausible that WNT5A functions chiefly within the dermis, reprising its role in condensate formation and directional outgrowth of structures like the limbs and phallus (*22*). In this case, epidermal hyperplasia may only be an incidental readout of a distinct mesenchymal signaling environment and might thus be interpreted as an evolutionary spandrel. Changes in the relative mechanical properties of the dermal and epidermal layers of the patagium primordium driven by WNT5A may, however, also play a distinct role in promoting formation of the initial patagium ridge, for instance through buckling morphogenesis (*44*). This process can occur when two adherent tissue layers expand at different rates, leading to a mechanical instability that forces the formation of a curvature. For instance, it has been proposed that gyrification in the brain may be caused by frustration of gray matter expansion due to its physical connection to underlying, slow-growing white matter (*45*). Similar tissue buckling has also been shown in the frilled dragon (*Chlamydosaurus kingii*). This species possesses several highly stereotyped convex folds on the skin constituting its eponymous neck frill which arise during embryogenesis through frustrated growth of its anterior surface (*46*). If related mechanisms are at play in flying mammals, WNT5A may drive localized stiffening of mesenchyme in the patagium primordium, thereby constraining the expansion of overlying epidermis and promoting a convex curvature of the lateral skin, like that seen in the early patagium ridge (**Fig. 1C and Fig. 4C**).

Interestingly, we find further evidence of molecular convergence between sugar gliders and bats. During the patagium outgrowth phase in both species, we discover many shared differentially expressed genes, including significant enrichment of genes with roles in limb development.

Because limbs are ancestral to all tetrapods, components of their underlying developmental programs may represent a molecular toolkit that diverse lineages may have drawn upon during the evolution of flight. Such a process may also be favored by evolution, as it could promote adaptive functional integration between the patagium and limbs. In both sugar gliders and bats, the patagia must be physically connected to the limbs in order to function. Yet, limbs and patagia do not arise simultaneously during development. Given that spatial patterning mechanisms already exist in tetrapods to specify outgrowth programs to the lateral plane during limb formation, these mechanisms may inherently provide a means to direct patagium growth to its proper location in relation to the limbs. Exploring similarities between the regulatory apparatuses of convergently re-deployed genes, such as *Wnt5a, Tbx5, Tbx3, Hand2*, and *Osr1* in the limbs and lateral patagia, presents a valuable future direction, as it has the potential to recalibrate our understanding of the role of pleiotropy in adaptive evolution. Most often, pleiotropy is viewed as an important source of evolutionary constraint, because recycling of developmental programs exposes them to greater negative selection (*47*). In the case of the patagium however, reuse of mechanisms for spatially patterning the expression of limb outgrowth genes in a single coronal plane may provide a means of facilitating beneficial functional integration between an ancestral structure and a novel one. Therefore, continued exploration of the gene networks governing patagium formation in mammals and their regulatory architecture has the potential to provide profound insights into development and adaptation.

## Acknowledgements

We thank Princeton LAR (Kirsten Gerhart, Grace Barnett, and Jamus McGuire) for help with sugar glider husbandry; the LSI Genomics Core (Jennifer M. Miller and Jean Arley Volmar) for help with library preparation and sequencing; Tom Pisano for help with tissue clearing and light-sheet imaging; Forrest Rogers for help with statistics; John Scheibe for access to initial sugar glider samples; and Elise Ireland for proofreading.

## Author contributions

RM and CF conceived the study and designed experiments. JC, RRB, and JJR provided bat tissues and BO provided initial sugar glider tissues. CF, AA, SM, JM, BO, and RM performed phenotypic characterization and histological analyses of sugar glider and bat tissues. RM generated the light-sheet movie. WW designed the sequencing strategy for sugar glider and bat transcriptomic samples and for the sugar glider genome. CF performed all computational analyses, including assembly and assessment of the sugar glider genome, generation of gene annotations for sugar glider and bat, design of sugar glider *in situ* hybridization probes, as well as the design and execution of all transcriptomic analyses in both the sugar glider and bat. CF conducted tissue recombination and bead implantation experiments in sugar glider explants. SM, RR, MP, and RM conducted mouse experiments and phenotypic characterizations. RvA generated *Wnt5a;R26rtTA* mice. RM, AA, and CF generated data visualizations. RM supervised the project and provided administration. RM and CF curated data generated in the study. CF and RM wrote the manuscript with input from all other authors.

## Data and materials availability

The sugar glider genome assembly, corresponding genome sequencing reads and RNA-Seq reads used in transcriptomic analyses are submitted under NCBI BioProject: PRJNA849992. RNA-Seq reads for Seba’s short-tailed bat are submitted under NCBI BioProject: PRJNA859189. Genome annotations for the sugar glider and Sebat’s short-tailed bat are available in the following FigShare repository: https://figshare.com/s/d6c585fbae0c1f22e8df. Code used for differential expression and WGCNA analyses are hosted on GitHub: https://github.com/charlesfeigin/Petaurus-glider-transcriptomics-code.

## Materials and Methods

### Sugar Glider Husbandry

Captive-born, adult sugar gliders were obtained from the US pet trade and thereafter were maintained in breeding colonies at Princeton University and Grand Valley State University. All experiments performed were approved by the respective IACUC committee at Princeton University and Grand Valley State University. The colonies were maintained on a 12hr:12hr light-dark cycle, within temperature and humidity rates of 20-27°C and 30-70% respectively. Animals were housed as bonded male-female pairs or groups of 1 male and 2-3 females and fed a diet consisting of dried food, fruits, and protein. Animals were checked for pouch young typically 2-5 times per week by gentle palpation of the maternal pouch and provided a drop of honey as a reward, delivered on a disposable tongue depressor. Pouch young discovered by palpation were visually examined by briefly anesthetizing the mother with isoflurane and gently everting the material pouch to expose the neonate. All pouch young were either found alone or as twins, with one on each side of the pouch. Pouch young, which remain attached to the nipple during the first several weeks of life, were removed while the mother was under anesthesia by gently placing the pointer finger and thumb in front of its snout and applying light pressure.

### Species Nomenclature

The marsupial gliders kept in the US pet trade are commonly referred to as sugar gliders and belong to the *Petaurus breviceps* species complex, whose native range extends through significant parts of eastern mainland Australia, as well as the island of New Guinea (both Papua New Guinea and West Papua, Indonesia) (*9*). Studies over the last decade indicate that genetic relationships of populations across this range are complex (*48-50*). However, findings by Campbell et al. 2019 suggest that gliders in the US pet trade, from which our sugar glider colony was derived, originated from a source population in West Papua Indonesia, near the city of Sorong and may have sister-group relationship to Australian populations (*50*).

Recently, Cremona et al. 2020 proposed that several Australian populations within the *P. breviceps* complex, be split into three distinct species: *P. breviceps* (sugar glider), *P. notatus* (Krefft’s glider) and *P. ariel* (savanna glider), occupying abutting ranges in eastern Australia. This proposed division of *P. breviceps* was based on highly-integrative, but fairly limited data (*49*). Importantly, this work did not address the complex relationship between New Guinean populations with those in Australia and no new nomenclature for New Guinean populations was put forward. Consequently, the proposed species division could, in light of the phylogeographic relationships suggested by (*50*), result in paraphyly of *P. breviceps*. Thus, while recent studies point to previously unappreciated genetic diversity and structure within these gliders that may ultimately warrant reorganization and/or division of the *P. breviceps* complex, determining reliable species boundaries will likely require further systematics research.

Given this, when referring to the complex of marsupial gliders to which our model organism belongs, we have opted to continue the use of *P. breviceps*. Additionally, following the convention of the pet trade from which our research colony was derived, we use the common name sugar glider. We recognize however that the taxonomy of *Petaurus* gliders is currently in flux and is likely more complex than current nomenclature reflects. Future population genomic studies of both Australian and New Guinean *Petaurus* gliders will thus be of tremendous interest to the systematics, evolution, and conservation research communities.

### Sugar Glider Genome Sequencing, Assembly and Annotation

#### Genome sequencing and assessment

50mg of muscle tissue were obtained from a euthanized adult female sugar glider and high-molecular weight DNA was extracted using the Qiagen MagAttract HMW kit (NEB 67563). DNA was quantified using the Qubit DNA HS kit (Thermofisher Q32851) and fragment size was assessed using Bioanalyzer 2100 and High Sensitivity chip (Agilent 5067). Sequencing libraries were prepared using the Genome v2 Library Prep kit and Chromium instrument (10X Genomics). Libraries were then sequenced on an Illumina HiSeq 2500 Rapid Flowcell to produce ∼241.5Gb of sequence in paired-end 2×200bp format. Reads were assembled using the SuperNova2.0 pipeline (10X Genomics) (*10*). A small number of duplicate scaffolds were identified by reciprocal blast (see below) and removed. Summary statistics for the resulting assembly were generated using the stats.sh script in the BBmap package (v37.93). Representation of mammalian universal single-copy orthologs in the sugar glider and other diprotodont marsupials were assessed with BUSCO version 5.5.2 (https://busco.ezlab.org/) using the curated mammalian version 10 database (mammalia_odb10) (*51*). Values for the common wombat (*Vombatus ursinus*), koala *(Phascolarctos cinereus*), tammar wallaby (*Macropus eugenii*), quokka (*Setonyx brachyurus*) and ground cuscus (*Phalanger gymnotis*) were generated previously (*52*). Values for all other species shown were calculated in the present study.

#### Gene annotations

Genes in the sugar glider assembly were annotated with the Maker2 pipeline using multiple lines of evidence (*11*). Homology evidence was generated through alignment of RefSeq proteins and transcripts sequences from koala (*Phascalarctos cinereus*), Tasmanian devil (*Sarophilus harrisii*), gray short-tailed opossum (*Monodelphis domestica*), tammar wallaby (*Notamacropus eugenii*), human (*Homo sapiens*) and laboratory mouse (*Mus musculus*) (*12, 14*). *De novo* sugar glider transcripts were assembled from RNA-Seq reads using Trinity v2.7.0 with default settings (*53*). Transcripts were aligned to the genome as EST evidence. *Ab initio* gene predictions were performed using the human model provided by Augustus and a sugar glider model generated using SNAP (*13, 54*). Three annotation iterations were performed in total. In the first, only the Augustus model and alignment evidence were used (sugar glider transcripts and homologous proteins/transcripts), with parameters est2genome and protein2genome set to 1. Gene models with an annotation edit distance (AED) < 0.75 were then used to generate a hidden Markov model (HMM) for SNAP. In the second iteration, the initial SNAP HMM was included along with all other of evidence, and est2genome and protein2genome were set to 0. A final SNAP HMM was then produced and included in a third and final iteration. Identities were assigned to gene models by reciprocal blast implemented by crb-blast (https://github.com/cboursnell/crb-blast) against Refseq koala transcripts (*12, 14*). Annotations for the sugar glider were deposited in a FigShare repository (https://figshare.com/s/d6c585fbae0c1f22e8df).

### Phenotypic characterization of sugar glider pouch young and bat embryos

#### Sample processing

Sugar gliders: Sugar glider pouch young were injected intraperitoneally with 50mg/kg of 5-ethynyl-2’-deoxyrudine (EdU; Lumiprobe) and were collected 2hrs later. Samples used for section histology were then fixed in 4% paraformaldehyde (PFA) and processed through the following overnight incubations: 10% sucrose-PBS, 30% sucrose-PBS and a 50:50 mixture of 30% sucrose-PBS and optimal cutting temperature (OCT) compound. Processed samples were then embedded in OCT (Tissue-Tek: 4583) and transverse sections 10-14μm were taken using a cryostat (Leica: CM 3050S). EdU incorporation was visualized by using a labeling mix containing Copper Sulfate Pentahydrate (Sigma: 209198, 2mM), Ascorbic Acid (Sigma: PHR1008, 20mg/mL), and Sulfo-Cyanine5 Azide Dye (Lumiprobe: A3330, 8μM). Samples were also washed for 10 minutes in a solution of 1.25×10^−3^ mg/mL of 4′,6-diamidino-2-phenylindole (DAPI; Sigma: D9542) in PBS to fluorescently label nuclei within tissue sections.

#### Bats

Bat embryos (*Carollia perspicillata*) were wild collected in Trinidad under permits issued by the Wildlife Section, Forestry Division, Ministry of Agriculture, Land and Marine Resources (Republic of Trinidad and Tobago). Embryos were fixed in 4% PFA in PBS for 24hrs, rinsed in PBS, and gradually dehydrated in increasing concentrations of methanol (25, 50, 75, 100% in doubly distilled water (ddH_2_O), 1hr each) and stored at -80°C. Upon arrival to the laboratory, samples were either processed for section histology and immunohistochemistry (IHC), as described above, or kept in methanol for whole-mount IHC.

#### Confocal imaging

Confocal stacks from sections of sugar glider pouch young were generated using a A1R-STED confocal microscope (Nikon) and processed with the accompanying NIS-Elements software (Nikon) Confocal stacks were then split into single channel stacks (488 and 647 respectively). The brightness of each stack was adjusted to maximize the signal-to-noise ratio and then stacks were flatted into single reference images with ImageJ’s built-in “Z Project” function. Binary conversion was done using ImageJ’s “Make Binary” function with the IsoData method and a dark background. Thresholds were calculated using the entire Z projection. The resulting images had two pixel values, 0 for black and 255 for white. All pictures are representative of three individuals per stage.

#### Quantification of epidermal thickness and proliferation

The epidermis was identified by eye using the DAPI channel in ImageJ and divided into 20 equal regions of interest (ROIs) along the dorsal-ventral axis. A segmented line was drawn following the curve of the basement membrane of the epidermis from the neural tube to the ventrum. This line was used to define 20 ROIs of equal lengths along the basement membrane, extending distally to the outer edge of the epidermis. Epidermal thickness was quantified by measuring the distance from the basement membrane to the exterior margin of the epidermis at the distal edge of each ROI. Epidermal proliferation was quantified in the EdU channel using ImageJ’s “Analyze Particles” function to identify dividing cells in each epidermal ROI. Pictures are representative of three individuals per stage.

#### Quantification of dermal density and proliferation

The dermis underlying each epidermal ROI was defined along the same segmented line following the curve of the epidermal basement membrane described above. A straight line 100 microns in length was drawn toward the center of the tissue section from the edge of each epidermal ROI. These lines were connected by another segmented line following the curvature of the tissue, resulting in a region of dermis underlying each of the previously defined epidermal ROIs. Dermal density was quantified in the DAPI channel using ImageJ’s “Analyze” function to find the mean fluorescence intensity (MFI) of the DAPI signal for each dermal ROI. This produced a measure of the average intensity of signal adjusted for the area of the ROI. Dermal proliferation was quantified in the EdU channel using ImageJ’s “Analyze Particles” function to identify dividing cells in each dermal ROI. Pictures are representative of three individuals per stage.

#### Tissue clearing and light sheet microscopy for sugar glider joeys

Postnatal day 3 sugar glider joeys were injected with EdU, collected 2hrs later, and fixed in 4%, as described above. Visualization of cell proliferation was carried using a modified iDISCO and tissue clearing protocol (*55, 56*). Briefly, joeys were dehydrated in increasing concentrations of methanol (25, 50, 75, 100% in doubly distilled water (ddH_2_O), 1hr each), bleached in a 5% hydrogen peroxide (H_2_O_2_; Sigma: 1086001000)/methanol solution overnight, and serially rehydrated (methanol:ddH_2_O, 1 hr each). Joeys were then washed in a solution of 20% DMSO (Fisher Scientific: D128) + 0.3M Glycine (Sigma: 410225) + 0.2% Triton X-100 (Sigma: T8787)/PBS at 37C for 2 days and subsequently immersed in a blocking solution of 10% DMSO + 6% donkey serum (EMD Millipore: S30) + 0.2% Triton X-100/PBS at 37C for 3 days. After this, joeys were washed twice for 1hr/wash in a solution containing 0.2% Tween-20 (Sigma: P9416) + 10mg/ml heparin (Sigma: H3149)/PBS. Following this, joeys were processed for EdU incorporation using the Click-iT EdU Imaging kit (Thermo Fisher: C10086), following manufacturer’s suggestions, except that incubation times were extended to 2 days for each step. After the final wash, samples were serially dehydrated in methanol (methanol:ddH_2_O, 1hr each), treated with 2:1 dichloromethane (DCM; Sigma: 270997):methanol and then incubated in 100% DCM for 2 days. Embryos were then placed in dibenzyl ether (DBE; Sigma: 108014) and stored at room temperature prior to imaging. For imaging, joeys were glued lateral side down on a custom-designed 3D printer holder and imaged in DBE using a light-sheet microscope (Ultramicroscope II, LaVision Biotec) running ImSpector Microscope controller software (version 5.1.347). Imaging of EdU+ cells was acquired as described previously (*56*).

### Spatial analysis of gene expression

#### Fluorescence in situ hybridization (FISH)

Starting from a pool of sugar glider skin cDNA, we generated DIG-labeled riboprobes by cloning fragments of *Wnt5a* (primers: 5’-GAGACGGCCTTCACTTATGC-3’ and 5’- GTCTGCACCGTCTTGAACTG-3’), *Wnt11* (primers: 5’-AACAGCTGGAAGGACTGGTG-3’ and 5’-CACCTTGGTGGCTGATAGGT-3’) and *Dkk2* (5’- CCCAGTTACCGAAAGCATATTAACC-3’ and 5’- AAAGCCAGACTCCATGTGTGTC-3’). We carried out FISH following previously described protocols (*57*). Briefly, tissue sections were post-fixed with 4% PFA in PBS, washed with PBS, placed in Triethanolamine/Acetic Anhydride for 20min, and incubated overnight with riboprobes at 65°C. The following morning, probes were washed with 0.2X Saline Sodium Citrate, equilibrated with buffer, and incubated with secondary anti-DIG antibody (1:500) for 3hrs. After washing several times, the signal was developed with the Tyramide Signal Amplification System, following the protocol suggested by the manufacturer (Perkin Elmer: NEL 753001KT). Tissues were visualized using an A1R-STED confocal microscope (Nikon). All pictures are representative of at least three individuals per stage.

#### Immunohistochemistry

Immunohistochemistry in sections of sugar glider explants, transgenic mouse skin and bat plagiopatagium was performed using anti-KRT14 (BioLegend: 905303; 1:1000), anti-WNT5A (BiossUSA: bs-1948R; 1:100), and anti-Ki67 (Abcam: ab15580; 1:100). Reactions were visualized with HRP-streptavidin and the AEC substrate kit (Vector Labs: SK4200) or Alexa-dye-conjugated secondary antibodies (Thermo Fisher). Control tissue was incubated with PBS instead of primary antibodies. Cell nuclei were stained with DAPI. Sections were examined using an A1R-STED confocal microscope and a NiE upright microscope (Nikon). All pictures are representative of at least three individuals per stage. Immunohistochemistry of the mouse pinna was performed using the rabbit monoclonal Cytokeratin 14 (K14) antisera (SP53, ab119695; abcam) for 8 hours at 4°C, followed by incubation with Goat Anti-Rabbit IgG H&L Alexa Fluor® 555 (ab150078; abcam) for 1 hour at room temperature.

#### Whole-mount in situ hybridization

Whole-mount *in situ* hybridization on bat embryos was carried out following a previously established protocol with slight modifications(*58*). Briefly, embryos were rehydrated in increasing concentrations of methanol (25, 50, 75, 100% in PBS, 20min each) and postfixed in a solution containing 4% PFA + 0.1% Glutaraldehyde (Sigma G5882). Embryos were then treated with 5% H_2_O_2_ for 1hr to block endogenous peroxidase activity, washed in PBT (1xPBS + 0.1% Tween-20), and incubated for 1hr in blocking solution (3% BSA (Sigma: B4287) + PBS + 0.1% Tween 20). Following this, we incubated embryos overnight with primary antibody diluted in blocking solution. The next morning, embryos were washed in PBS for 1hr, treated with 3% H_2_O_2_ for 1hr, and washed with PBT for 1hr. Embryos were then incubated overnight with secondary biotinylated antibody (Vector Labs: BP-9100-50). The following day, embryos were washed and incubated with HRP-Streptavidin (Vector Labs: SA-5704-100) for 1hr and developed the signal with AEC (Vector Labs: SK4200, following the manufacturer’s instructions.

#### RNA scope

Cells expressing *Wnt5a* were visualized in the pinnae of E14 embryonic mice from cryosectioned tissues (12μm sections) using the RNAscope® v2 (323100, Advanced Cell Diagnostics), following the manufacturer’s instructions. The *Mus musculus Wnt5a* probe (316791; Advanced Cell Diagnostics) was used. For all experiments, slides were mounted using VECTASHIELD® Antifade Mounting Medium and with DAPI (H-1200-10; Vector Laboratories). Fluorescent images were captured with a Fluoview3000 laser-scanning microscope (Olympus) with a UPLSAPO 10X/0.45 and 40X/1.25 Sil objectives (Olympus).

### RNA sequencing and data preprocessing

#### Sugar gliders

We collected 34 patagium samples from glider pouch young spanning the first three weeks of postnatal development, as well as dorsal skin from approximately the first 12 days postnatal and shoulder skin from approximately the first 4 days postnatal. RNA was extracted from each tissue by first lysing skin tissue with the Next Advanced Bullet Blender and then isolating total RNA with the RNeasy Fibrous Tissue Mini Kit (Qiagen: 74704). 100ng of RNA was provided to the Genomics Core Facility at the Lewis-Sigler Institute (Princeton University) for quality control and library preparation following the Smart-seq2 protocol with an average insert size of 200bp (*59*). Size distribution and RNA integrity were measured using the Agilent Bioanalyzer 2100 and RNA 6000 Pico chip. Libraries were individually barcoded, pooled and then sequenced at the New York Genome Center on an Illumina NovaSeq in 2×150bp format. After demultiplexing, reads were filtered by quality and trimmed to remove residual adapter sequences and low-quality end-bases using Trimmomatic 0.38 parameters: SLIDINGWINDOW:5:20 MINLEN:50 AVGQUAL:25) (*60*). Sequenced RNA libraries were deposited under BioProject: PRJNA849992, BioSample: SAMN29758252. Reads were then mapped against the sugar glider genome assembly using STAR-2.7.3a (*61*).

#### Bats

Bat embryos were wild collected and preserved in RNAlater (ThermoFisher Scientific). Upon return to the lab, dorsal (N = 5) and lateral skin tissues (N = 5) were dissected. RNA was extracted and its quality assessed following the same procedures outlined above. Libraries were prepared using the directional PrepX RNA-seq library preparation protocol on the automated Apollo 324TM NGS Library Prep System (Takara Bio, CA), with a cDNA insert average size of ∼200bp.

To produce genome annotations suitable for RNA-Seq analyses, we performed an annotation lift-over procedure implemented in the program Liftoff v1.6.3 (parameter: -d4) to transfer gene models from the closely-related microbat *Artibeus jamaicensis* (GCF_014825515.1/WHU_Ajam_v2) to the DNAZoo *Carollia perspicillata* assembly (*62, 63*). This yielded 23,706 gene models used for downstream analysis. Annotations for Seba’s short-tailed bat were deposited in a FigShare repository (https://figshare.com/s/d6c585fbae0c1f22e8df).

#### Weighted Gene Correlation Network Analysis

Network analyses were performed using Weighted Gene Correlation Network Analysis (WGCNA) (*15*). Briefly, read counts generated by featureCounts were normalized by rlog transformation (*17*). The pickSoftThreshold function was used to estimate scale-free threshold for each dataset, testing soft power thresholds from 1 to 30 in increments of 1. We chose a scale-free topology of 0.8 (80%), which was achieved at a soft power threshold of 11. An unsigned correlation network was them constructed with the blockwiseModules function using the soft power threshold determined above. Gene correlation calculations are agnostic of sample stage or weight. Therefore, after module construction samples were ordered by developmental stage and animal weight, then per-sample module eigengene values of each module were plotted as a heatmap (**Fig. 2A**). The correlation coefficient (weighted Pearson correlation) between PM8 module eigengene expression and animal weight (a proxy for age) was calculated and its significance assessed by Student’s *t*-test using WGCNA’s included cor and corPvalueStudent functions respectively (*15*). PM8 genes were examined for gene ontology term enrichment via the Gene Ontology Resource web server (http://geneontology.org/), using the *Mus musculus* biological process database. Term enrichment significance was determined by Fisher’s exact test (FDR < 0.05).

#### Differential Gene Expression Analyses

Pairwise differential expression analyses between the transcriptomes of stage-matched patagium, dorsal and shoulder skin from the sugar glider or plagiopatagium and dorsal skin from Seba’s short-tailed bat were performed using DeSeq2 v1.30.1 from BioConductor (https://bioconductor.org/) (*17*). For each analysis, the dataset was re-leveled such that the non-patagium comparison tissue (i.e., dorsal skin or shoulder) was set as the reference level so that patagium-upregulated genes were expressed as positive log2fold changes and patagium-downregulated genes were negative. The DESeq2 function was called and resulting p-values were conservatively adjusted for multiple testing using the Benjamini-Hochberg procedure (FDR < 0.01) and only genes showing at least a 1.5-fold change were considered differentially expressed. Gene Ontology analyses for genes upregulated in the sugar glider lateral patagium or bat plagiopatagium were performed using the Gene Ontology Resource web server (http://geneontology.org/) and the *Mus musculus* biological process database. Term enrichment significance was determined by Fisher’s exact test (FDR < 0.05).

### Epidermal-dermal recombination explant cultures

Pouch young were collected at approximately P4. This stage was used as it represents the period prior to patagium outgrowth when dermal condensation becomes most defined and because enzymatic dissociation of live dermis and epidermis prior to this stage had a low rate of success. For each pouch young, patagium skin and dorsal skin were removed using fine dissecting scissors and ultrafine forceps. Each tissue was then floated in one well of a 6-well plate containing 3mL of DMEM (Corning: 10-017-CV) plus 0.3g dispase II (Sigma: D4693) with the epidermis facing upward and the dermis submerged in media. Tissues were then incubated on a nutator rotating at 12rpm for 2 hours at 37°C. At the 1 hour and 1.5 hour marks, toothbrush bristles were used to gently tease apart epidermis and dermis to expose new surface area to the enzyme. After dissociation was complete, tissues were transferred to wells of a fresh six-well plate with 3mL DMEM + 10% Bovine Calf Serum (Cytiva HyClone:SH3007203; hereafter BCS) and nutated at 12rpm for 15 minutes at room temperature to wash away residual enzyme. A single Whatman Nucleopore Track-Etched Membrane (WHA110614) was placed in the lid of a sterile petri dish, and fine forceps were used to transfer a dermis (either patagium or dorsal control) into the dish. Forceps were used to slide the dermis onto flat paddle forceps (pre-wetted in DMEM + 10% BCS). The dermis was then gently slid onto the membrane. The wetted paddle forceps were then used to lift segments of dorsal epidermis from the surface of the wash media, and toothbrush bristles were used to spread it on top of the dermis. The whole membrane, recombined explant in place, was then transferred to a fresh six-well plate with 3mL of DMEM + 10% BCS + 1**%** penicillin/streptomycin (Gibco: 15070-063). Two recombinations were performed, each with 5 replicates. Dorsal epidermis was recombined with patagium dermis to test the potential for patagium dermis to drive morphological changes in epidermis. As a negative control, dorsal epidermises were recombined with dorsal dermis. Explants were cultured for 96hrs, replacing media each day. Explants were then fixed while attached to the membrane filter in 4% PFA for approximately 4 hours, before being processed (undergoing overnight incubations in 10% sucrose, 30% sucrose, 50:50 mix of 30% sucrose and OCT) and embedded in OCT. Sectioning, was performed as described above. Epidermal thickness was quantified using Fiji software by measuring tissue stained with Hematoxylin and Eosin from the base of the epidermis to the top of the epidermis. We took a total of 10 measurements/tissue section, 3 tissue sections/sample, 5 samples/treatment. Statistical significance was assessed using a general mixed effects model ANOVA test (fixed effect = treatment; random effect = individual/sample). Statistical significance was assigned to p-values: p<0.05(*); p<0.01(**), p<0.001(***), p<0.0001(****).

### Bead implantation in dorsal skin explants

100uL of Bio-Rad Affi-gel blue (Bio-Rad: 1537301) were added to 1mL sterile PBS in a 2mL low-bind tube (Eppendorf: 0030108450), vortexed briefly, then centrifuged for 10 seconds in a microfuge. PBS was removed by pipetting and the procedure was repeated two more times. Beads were then resuspended in a final volume of approximately 10-20uL. Concentrated beads were then placed in a sterile petri dish, divided into two equal sized droplets and allowed to dry in a sterile hood for at least 2 hrs. When beads were dry, they were resuspended in a protein solution. To test *Wnt5a* function, beads were soaked in 2uL of a 50:50 mix of recombinant WNT5A (R&D Systems: 645-WN-010) at stock concentration of 1ug/uL and PBS with 50ug/uL BSA. For controls, beads were soaked in a 25ug/uL BSA. Beads were incubated in their respective solutions at 4 °C for 1.5hrs, during which time pouch young at approximately P4 were collected. Pouch young were euthanized 5-10 minutes before the end of bead incubation and placed under a dissecting scope. The bevel of a 31-gauge needle was then used to create several pockets into the dermis of the dorsal skin, into which beads were placed using sterile #55 forceps. After beads were placed, the dorsal skin was removed from the pouch young and placed onto Whatman filter membranes (Whatman: 131135012e) using forceps and toothbrush bristles and grown for 72hrs as explants floating on full media DMEM + 10% BCS + 1**%** penicillin/streptomycin. Explants were then fixed while attached to the membrane filter in 4% PFA for 3hrs, before being processed for embedding and sectioning, as described above. Epidermal thickness was quantified using Fiji software by measuring tissue stained with Hematoxylin and Eosin from the base of the epidermis to the top of the epidermis. We took a total of 10 measurements/tissue section, 3 tissue sections/sample, 5 samples/treatment. The first two measurements were taken directly above the bead, with subsequent ones taken in opposite directions (4 to the left of the bead and 4 to the right, equidistantly spaced). Statistical significance was assessed using a general mixed effects model ANOVA test (fixed effect = treatment; random effect = individual/sample). Statistical significance was assigned to p-values: p<0.05(*); p<0.01(**), p<0.001(***), p<0.0001(****).

### Mouse experiments

#### Mouse strains

The following mouse strains were obtained from Jackson laboratories: FVB/N-Tg(tetO-Wnt5a)17Rva/J (JAX: 022938); and B6.Cg *Gt(ROSA)26Sor*^*tm1(rtTA*M2)Jae*^/J (JAX: 006965).

#### Induction of transgenes and tissue processing

For *Wnt5a* induction, P30 mice were placed on 1mg/ml Doxycycline containing water *ad libitum* for 7 days (Doxycycline water was replaced every other day). Tissues were fixed and processed for histology and IHC, as described for the sugar gliders and bats.

#### Phenotypic characterization

*Wnt5a* transgenics: Epidermal thickness was quantified using Fiji software by measuring tissue stained with Hematoxylin and Eosin from the base of the epidermis to the top of the epidermis. Measurements were taken exclusively from interfollicular regions (10 measurements/dorsal skin section, 3 dorsal skins sections/sample, 4 samples/treatment). Cell density was quantified on tissue sections by counting the number of DAPI+ cells per surface area (measurements were done on 3 dorsal skin sections/sample, 4 samples/treatment). Statistical significance was assessed using a general mixed effects model ANOVA test (fixed effect = treatment; random effect = individual/sample). Statistical significance was assigned to p-values: p<0.05(*); p<0.01(**), p<0.001(***), p<0.0001(****).

#### Ear pinna

Quantification of cell nuclei and epidermal thickness was done using the ImageJ software. Briefly, we counted all DAPI-stained cell nuclei in K14+ epidermis across different zones of the embryo head, including ear pinna. Epidermal thickness was measured using the measure tool to measure thickness of the K14+ layer across different zones of the embryo head (20 measurements/zone across the epidermis, 3 embryos). Statistical significance was assessed using a general mixed effects model ANOVA test (fixed effect = treatment; random effect = individual/sample). Statistical significance was assigned to p-values: p<0.05(*); p<0.01(**), p<0.001(***), p<0.0001(****).

