## Supplementary Figures and Tables for "Convergent deployment of ancestral programs during the evolution of mammalian flight membranes"

**Fig. S1.**

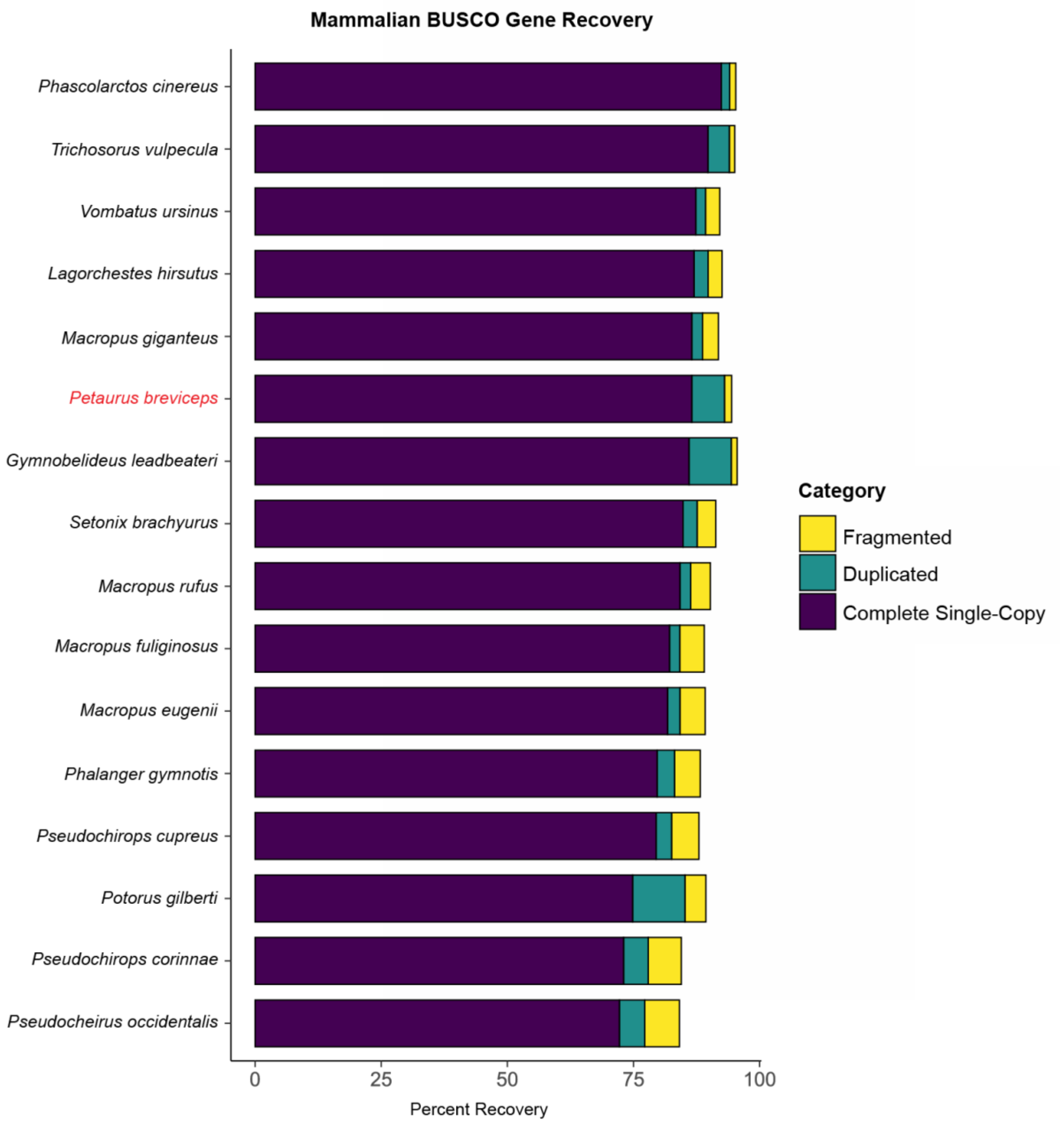


**Comparison of benchmarking universal single-copy ortholog (BUSCO) recovery in the genomes of the sugar glider and related diprotodont marsupials.** Species are sorted by the percent of complete-single copy BUSCO genes. The sugar glider assembly presented in this study is shown in red font.

**Fig. S2.**

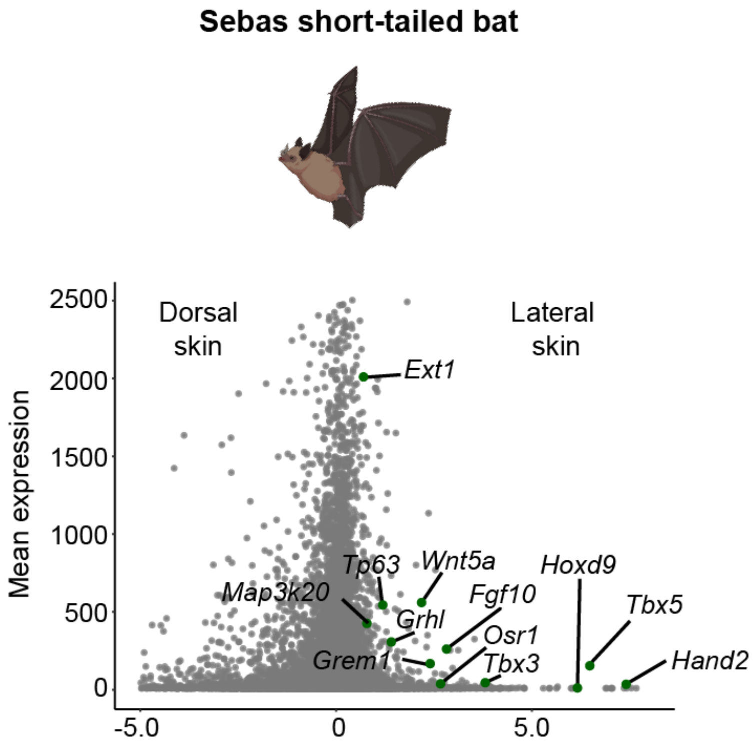


**Volcano plot of genes differentially expressed between the bat plagiopatagium and neighboring dorsal skin during its outgrowth.** Genes upregulated in the plagiopatagium and belonging to limb development Gene Ontology terms are labeled in green.

**Table S1.**
Sugar glider genome assembly metrics.

| Metric | Value |
|---|---|
| Scaffold Length | 3463.91 Mbp |
| Gap-Free Length | 3344.58 Mbp |
| Contig/Scaffold Number | 87,817 / 1,909 |
| Contig/Scaffold N50 | 69.63 Kbp / 28.19 Mbp |
| Contig/Scaffold N90 | 51.92 Kbp / 20.28 Mbp |
| Longest Contig/Scaffold | 599.68 Kbp / 109.60 Mbp |
| Gap Percentage | 1.89% |
| G+C Percentage | 38.95% |

**Table S2.**
Sugar glider genome BUSCO recovery.

| **BUSCO Reference Database: mammalia_odb10: 9226 genes** | | |
|---|---|---|
| **BUSCO category** | **Gene Count** | **Percent Recovery (of 9226)** |
| Complete (total) | 8581 | 93.0 |
| Complete single-copy | 7984 | 86.5 |
| Complete and duplicated | 597 | 6.5 |
| Fragmented | 132 | 1.4 |
| Missing | 513 | 5.6 |

**Table S3.**
Manifest of RNA-Seq samples used in this study.

| **Species** | **Sample** | **Tissue** | **Stage** | **Weight** |
|---|---|---|---|---|
| *P. breviceps* | pb_pat_pre-patagium_121 | Patagium | pre-patagium | 0.22 |
| *P. breviceps* | pb_pat_pre-patagium_122 | Patagium | pre-patagium | 0.22 |
| *P. breviceps* | pb_pat_pre-patagium_130 | Patagium | pre-patagium | 0.23 |
| *P. breviceps* | pb_pat_pre-patagium_131 | Patagium | pre-patagium | 0.25 |
| *P. breviceps* | pb_pat_pre-patagium_100 | Patagium | pre-patagium | 0.25 |
| *P. breviceps* | pb_pat_pre-patagium_137 | Patagium | pre-patagium | 0.26 |
| *P. breviceps* | pb_pat_pre-patagium_101 | Patagium | pre-patagium | 0.3 |
| *P. breviceps* | pb_pat_pre-patagium_119 | Patagium | pre-patagium | 0.35 |
| *P. breviceps* | pb_pat_pre-patagium_138 | Patagium | pre-patagium | 0.35 |
| *P. breviceps* | pb_pat_pre-patagium_118 | Patagium | pre-patagium | 0.35 |
| *P. breviceps* | pb_pat_early-forming_124 | Patagium | early-forming | 0.38 |
| *P. breviceps* | pb_pat_early-forming_129 | Patagium | early-forming | 0.39 |
| *P. breviceps* | pb_pat_early-forming_141 | Patagium | early-forming | 0.4 |
| *P. breviceps* | pb_pat_early-forming_132 | Patagium | early-forming | 0.41 |
| *P. breviceps* | pb_pat_early-forming_140 | Patagium | early-forming | 0.43 |
| *P. breviceps* | pb_pat_early-forming_139 | Patagium | early-forming | 0.49 |
| *P. breviceps* | pb_pat_early-forming_110 | Patagium | early-forming | 0.49 |
| *P. breviceps* | pb_pat_early-forming_102 | Patagium | early-forming | 0.52 |
| *P. breviceps* | pb_pat_early-forming_90 | Patagium | early-forming | 0.53 |
| *P. breviceps* | pb_pat_early-forming_103 | Patagium | early-forming | 0.55 |
| *P. breviceps* | pb_pat_early-forming_111 | Patagium | early-forming | 0.58 |
| *P. breviceps* | pb_pat_early-forming_142 | Patagium | early-forming | 0.59 |
| *P. breviceps* | pb_pat_early-forming_117 | Patagium | early-forming | 0.62 |
| *P. breviceps* | pb_pat_early-forming_105 | Patagium | early-forming | 0.63 |
| *P. breviceps* | pb_pat_early-forming_125 | Patagium | early-forming | 0.63 |
| *P. breviceps* | pb_pat_early-forming_120 | Patagium | early-forming | 0.69 |
| *P. breviceps* | pb_pat_late-forming_97 | Patagium | late-forming | 0.86 |
| *P. breviceps* | pb_pat_late-forming_93 | Patagium | late-forming | 0.89 |
| *P. breviceps* | pb_pat_late-forming_94 | Patagium | late-forming | 0.9 |
| *P. breviceps* | pb_pat_late-forming_108 | Patagium | late-forming | 0.9 |
| *P. breviceps* | pb_pat_late-forming_127 | Patagium | late-forming | 0.92 |
| *P. breviceps* | pb_pat_late-forming_126 | Patagium | late-forming | 0.97 |
| *P. breviceps* | pb_pat_late-forming_115 | Patagium | late-forming | 1.19 |
| *P. breviceps* | pb_pat_late-forming_98 | Patagium | late-forming | 1.68 |
| *P. breviceps* | pb_dors_pre-patagium_131 | Dorsal Skin | pre-patagium | 0.25 |
| *P. breviceps* | pb_dors_pre-patagium_137 | Dorsal Skin | pre-patagium | 0.26 |

| | | | | |
|---|---|---|---|---|
| *P. breviceps* | pb_dors_pre-patagium_101 | Dorsal Skin | pre-patagium | 0.3 |
| *P. breviceps* | pb_dors_pre-patagium_119 | Dorsal Skin | pre-patagium | 0.35 |
| *P. breviceps* | pb_dors_pre-patagium_138 | Dorsal Skin | pre-patagium | 0.35 |
| *P. breviceps* | pb_dors_pre-patagium_118 | Dorsal Skin | pre-patagium | 0.35 |
| *P. breviceps* | pb_dors_early-forming_124 | Dorsal Skin | early-forming | 0.38 |
| *P. breviceps* | pb_dors_early-forming_129 | Dorsal Skin | early-forming | 0.39 |
| *P. breviceps* | pb_dors_early-forming_141 | Dorsal Skin | early-forming | 0.4 |
| *P. breviceps* | pb_dors_early-forming_132 | Dorsal Skin | early-forming | 0.41 |
| *P. breviceps* | pb_dors_early-forming_140 | Dorsal Skin | early-forming | 0.43 |
| *P. breviceps* | pb_dors_early-forming_139 | Dorsal Skin | early-forming | 0.49 |
| *P. breviceps* | sg_shd_pre-patagium_131 | Shoulder Skin | pre-patagium | 0.25 |
| *P. breviceps* | sg_shd_pre-patagium_137 | Shoulder Skin | pre-patagium | 0.26 |
| *P. breviceps* | sg_shd_pre-patagium_101 | Shoulder Skin | pre-patagium | 0.3 |
| *P. breviceps* | sg_shd_pre-patagium_119 | Shoulder Skin | pre-patagium | 0.35 |
| *P. breviceps* | sg_shd_pre-patagium_138 | Shoulder Skin | pre-patagium | 0.35 |
| *P. breviceps* | sg_shd_pre-patagium_118 | Shoulder Skin | pre-patagium | 0.35 |
| *P. breviceps* | sg_shd_early-forming_124 | Shoulder Skin | early-forming | 0.38 |
| *P. breviceps* | sg_shd_early-forming_129 | Shoulder Skin | early-forming | 0.39 |
| *P. breviceps* | sg_shd_early-forming_141 | Shoulder Skin | early-forming | 0.4 |
| *P. breviceps* | sg_shd_early-forming_132 | Shoulder Skin | early-forming | 0.41 |
| *P. breviceps* | sg_shd_early-forming_140 | Shoulder Skin | early-forming | 0.43 |
| *P. breviceps* | sg_shd_early-forming_139 | Shoulder Skin | early-forming | 0.49 |
| *C. perspicillata* | sstb_pat_3_stg16 | Plagiopatagium | Stage16 | N/A |
| *C. perspicillata* | sstb_pat_5_stg16 | Plagiopatagium | Stage16 | N/A |
| *C. perspicillata* | sstb_pat_2_stg17 | Plagiopatagium | Stage17 | N/A |
| *C. perspicillata* | sstb_pat_4_stg17 | Plagiopatagium | Stage17 | N/A |
| *C. perspicillata* | sstb_pat_1_stg18 | Plagiopatagium | Stage18 | N/A |
| *C. perspicillata* | sstb_dors_stg16_3 | Dorsal Skin | Stage16 | N/A |
| *C. perspicillata* | sstb_dors_stg16_5 | Dorsal Skin | Stage16 | N/A |
| *C. perspicillata* | sstb_dors_stg17_2 | Dorsal Skin | Stage17 | N/A |
| *C. perspicillata* | sstb_dors_stg17_4 | Dorsal Skin | Stage17 | N/A |
| *C. perspicillata* | sstb_dors_stg18_1 | Dorsal Skin | Stage18 | N/A |

**Table S4.**

Gene models belonging to sugar glider patagium module 8.

| Gene Model ID |
|---|
| LOC110202101\|XM_020978119.1\|fragGeneModel-103-gene5-frag1 |
| PTBP1\|XM_021008785.1\|fragGeneModel-103-gene58-frag1 |
| MISP\|XM_021009057.1\|fragGeneModel-103-gene89-frag1 |
| TMEM59L\|XM_021009095.1\|maker-108-augustus-gene-3.21 |
| maker-108-snap-gene-8.17 |
| LOC110201946\|XM_020977849.1\|fragGeneModel-115-gene6-frag2 |
| WDR86\|XM_020977799.1\|maker-115-augustus-gene-9.8 |
| CNPY1\|XM_021009623.1\|maker-115-augustus-gene-27.9 |
| TEKT5\|XM_020972536.1\|fragGeneModel-118-gene11-frag1 |
| MAMDC2\|XM_021001775.1\|fragGeneModel-118-gene53-frag1 |
| NAAA\|XM_020988459.1\|maker-126-snap-gene-22.23 |
| SCD5\|XM_020988410.1\|maker-126-snap-gene-26.21 |
| SPP1\|XM_020988577.1\|maker-126-snap-gene-53.18 |
| SLC39A8\|XM_020988298.1\|maker-126-snap-gene-60.12 |
| TSPAN5\|XM_021008377.1\|maker-126-snap-gene-72.7 |
| UNC5C\|XM_021008379.1\|maker-126-augustus-gene-84.6 |
| TBCK\|XM_020974730.1\|maker-126-augustus-gene-114.9 |
| DKK2\|XM_021001334.1\|maker-126-augustus-gene-114.11 |
| OSTC\|XM_021001394.1\|maker-126-augustus-gene-123.7 |
| COL25A1\|XM_021001422.1\|fragGeneModel-126-gene47-frag2 |
| ENPEP\|XM_021001403.1\|maker-126-snap-gene-127.17 |
| PITX2\|XM_021001336.1\|fragGeneModel-126-gene41-frag2 |
| ARSJ\|XM_021001436.1\|snap_masked-126-processed-gene-141.5 |
| TENM3\|XM_021007594.1\|fragGeneModel-126-gene51-frag1 |
| NEIL3\|XM_020997850.1\|maker-126-snap-gene-213.15 |
| PALLD\|XM_020997845.1\|fragGeneModel-126-gene34-frag1 |
| KLHL2\|XM_020976334.1\|fragGeneModel-126-gene13-frag1 |
| SMAD4\|XM_020972302.1\|fragGeneModel-131-gene3-frag1 |
| TEX10\|XM_020995536.1\|maker-138-snap-gene-45.17 |
| MSANTD3\|XM_020995576.1\|maker-138-snap-gene-45.14 |
| ELP1\|XM_020995507.1\|maker-138-augustus-gene-46.7 |
| NMRAL1\|XM_020995621.1\|augustus_masked-138-processed-gene-60.1 |
| LOC110222032\|XM_021006845.1\|maker-138-augustus-gene-73.16 |
| METTL26\|XM_021006860.1\|maker-138-augustus-gene-77.21 |
| NARFL\|XM_021006659.1\|augustus_masked-138-processed-gene-78.10 |

| |
|---|
| MSLN\|XM_021006851.1\|maker-138-snap-gene-79.40 |
| LOC110221851\|XM_021006513.1\|maker-138-snap-gene-98.51 |
| maker-138-augustus-gene-101.65 |
| LOC110222072\|XM_021006913.1\|maker-138-augustus-gene-102.38 |
| TMSB10\|XM_020969493.1\|maker-139-snap-gene-2.26 |
| TRABD2A\|XM_020969508.1\|maker-139-augustus-gene-2.20 |
| CUNH20orf27\|XM_021001044.1\|maker-139-augustus-gene-31.9 |
| LOC110218385\|XM_021001060.1\|maker-139-snap-gene-45.23 |
| ADRA1A\|XM_021007717.1\|maker-139-snap-gene-60.4 |
| PCDH7\|XM_020992170.1\|maker-140-augustus-gene-50.12 |
| SLIT2\|XM_020992612.1\|fragGeneModel-140-gene26-frag1 |
| LAP3\|XM_020992719.1\|maker-140-augustus-gene-85.6 |
| PROM1\|XM_020992810.1\|maker-140-augustus-gene-88.11 |
| CYTL1\|XM_020992293.1\|maker-140-augustus-gene-113.6 |
| SORCS2\|XM_020992120.1\|maker-140-augustus-gene-127.4 |
| FGFRL1\|XM_021005458.1\|maker-140-augustus-gene-158.11 |
| TRIP13\|XM_020985654.1\|maker-151-augustus-gene-62.11 |
| UBE2QL1\|XM_021004424.1\|snap_masked-151-processed-gene-105.8 |
| ADCY2\|XM_021004443.1\|fragGeneModel-151-gene34-frag1 |
| CMBL\|XM_021004412.1\|maker-151-augustus-gene-115.13 |
| CCT2\|XM_020990239.1\|fragGeneModel-154-gene105-frag1 |
| IRX6\|XM_020984923.1\|maker-171-snap-gene-6.16 |
| PHKB\|XM_020985101.1\|maker-171-snap-gene-37.5 |
| ORC6\|XM_020985089.1\|maker-171-augustus-gene-39.16 |
| TSHZ3\|XM_020985057.1\|maker-171-snap-gene-55.9 |
| maker-171-augustus-gene-55.7 |
| PBX3\|XM_020967606.1\|fragGeneModel-171-gene9-frag1 |
| NEK6\|XM_020967527.1\|maker-171-augustus-gene-89.13 |
| NRTN\|XM_020971452.1\|maker-172-augustus-gene-0.11 |
| PODNL1\|XM_020994831.1\|maker-172-augustus-gene-9.20 |
| LYL1\|XM_020995079.1\|maker-172-augustus-gene-11.32 |
| MAST1\|XM_020994952.1\|maker-172-snap-gene-12.74 |
| ASNA1\|XM_020995306.1\|maker-172-augustus-gene-13.54 |
| PGLS\|XM_020994857.1\|fragGeneModel-172-gene11-frag1 |
| ZZEF1\|XM_020964666.1\|maker-173-snap-gene-3.37 |
| HES1\|XM_020964778.1\|fragGeneModel-173-gene2-frag1 |
| maker-173-snap-gene-38.50 |
| RARRES1\|XM_020988872.1\|maker-173-augustus-gene-62.37 |
| ECT2\|XM_020980475.1\|fragGeneModel-173-gene8-frag1 |

| |
|---|
| VOPP1\|XM_020981280.1\|maker-174-augustus-gene-35.9 |
| BARX1\|XM_020980965.1\|maker-174-augustus-gene-42.12 |
| TIMP4\|XM_020981072.1\|maker-174-augustus-gene-75.6 |
| FANCD2\|XM_020981005.1\|maker-174-augustus-gene-84.30 |
| ITPR1\|XM_020968454.1\|fragGeneModel-174-gene7-frag1 |
| LRRN1\|XM_020968476.1\|augustus_masked-174-processed-gene-107.0 |
| CNTN3\|XM_021007573.1\|fragGeneModel-174-gene60-frag1 |
| LOC110197030\|XM_020970590.1\|maker-174-snap-gene-211.18 |
| WNT5A\|XM_020970602.1\|maker-174-augustus-gene-221.10 |
| LRTM1\|XM_020970619.1\|maker-174-augustus-gene-223.4 |
| CACNA2D3\|XM_020970614.1\|fragGeneModel-174-gene19-frag1 |
| maker-174-snap-gene-237.14 |
| UBP1\|XM_020978524.1\|maker-174-augustus-gene-261.29 |
| NUF2\|XM_020978578.1\|maker-174-augustus-gene-262.41 |
| LOC110223842\|XM_021009534.1\|maker-175-augustus-gene-5.7 |
| ANXA1\|XM_021009517.1\|maker-175-augustus-gene-5.8 |
| ACTB\|XM_021008442.1\|maker-175-snap-gene-30.6 |
| SLC29A4\|XM_021008620.1\|augustus_masked-175-processed-gene-32.0 |
| AMZ1\|XM_020975264.1\|maker-175-snap-gene-44.17 |
| CUNH7orf50\|XM_020978697.1\|maker-175-snap-gene-58.23 |
| EEF2K\|XM_020965327.1\|maker-175-augustus-gene-83.12 |
| MRPS27\|XM_020998373.1\|fragGeneModel-175-gene28-frag1 |
| FAM172A\|XM_021008513.1\|maker-175-snap-gene-167.17 |
| ZFHX3\|XM_020963589.1\|fragGeneModel-176-gene2-frag1 |
| DPEP1\|XM_020989117.1\|maker-176-augustus-gene-14.33 |
| COTL1\|XM_020989254.1\|maker-176-augustus-gene-44.1 |
| snap_masked-176-processed-gene-44.0 |
| RRAD\|XM_020989437.1\|maker-176-augustus-gene-59.21 |
| LOC110193229\|XM_020964889.1\|maker-176-snap-gene-104.12 |
| UNC79\|XM_021000359.1\|snap_masked-177-processed-gene-15.4 |
| CRTAP\|XM_020987403.1\|maker-178-augustus-gene-2.16 |
| STEAP4\|XM_020987460.1\|maker-178-augustus-gene-19.4 |
| ITGA8\|XM_020987504.1\|maker-178-snap-gene-24.19 |
| DNAJC1\|XM_020966879.1\|fragGeneModel-178-gene5-frag2 |
| maker-178-augustus-gene-50.10 |
| maker-178-snap-gene-74.14 |
| CCNY\|XM_020990387.1\|fragGeneModel-178-gene27-frag1 |
| maker-178-snap-gene-114.32 |
| SDC2\|XM_020968874.1\|maker-179-augustus-gene-7.11 |

| |
|---|
| RAD54B\|XM_020968529.1\|maker-179-snap-gene-17.12 |
| CALB1\|XM_020979722.1\|maker-179-augustus-gene-34.9 |
| FSD1\|XM_020971283.1\|maker-180-snap-gene-7.32 |
| COQ9\|XM_020965528.1\|fragGeneModel-180-gene1-frag1 |
| AASS\|XM_020966361.1\|maker-181-augustus-gene-12.10 |
| ARF5\|XM_021002372.1\|maker-181-augustus-gene-36.15 |
| maker-181-snap-gene-39.41 |
| CCDC136\|XM_021002397.1\|augustus_masked-181-processed-gene-39.1 |
| COPG2\|XM_021002375.1\|maker-181-snap-gene-45.20 |
| PLXNA4\|XM_021002332.1\|fragGeneModel-181-gene28-frag1 |
| maker-181-snap-gene-59.5 |
| LOC110219169\|XM_021002303.1\|maker-181-augustus-gene-62.9 |
| TMEM139\|XM_020997927.1\|maker-181-snap-gene-92.43 |
| maker-182-snap-gene-5.23 |
| GINS3\|XM_020963767.1\|maker-182-snap-gene-11.19 |
| LOC110192444\|XM_020963594.1\|maker-182-augustus-gene-16.34 |
| CDO1\|XM_020967392.1\|maker-183-augustus-gene-45.14 |
| BAG1\|XM_020967377.1\|fragGeneModel-183-gene7-frag1 |
| DAPK1\|XM_020971674.1\|maker-183-augustus-gene-64.9 |
| FANCC\|XM_020971701.1\|maker-183-snap-gene-68.8 |
| MLLT3\|XM_021004579.1\|maker-184-snap-gene-6.3 |
| MEGF10\|XM_021009594.1\|maker-184-snap-gene-54.15 |
| CCK\|XM_020987376.1\|maker-185-augustus-gene-3.6 |
| PLCL2\|XM_020987210.1\|maker-185-snap-gene-31.9 |
| LRRC3B\|XM_020993310.1\|snap_masked-185-processed-gene-63.4 |
| MTURN\|XM_020993309.1\|maker-185-augustus-gene-86.9 |
| FKBP14\|XM_020993365.1\|maker-185-augustus-gene-87.16 |
| maker-185-augustus-gene-106.19 |
| GPC1\|XM_021005584.1\|maker-186-augustus-gene-15.11 |
| ABHD11\|XM_021005799.1\|maker-186-augustus-gene-28.17 |
| PSPH\|XM_021006954.1\|augustus_masked-186-processed-gene-49.5 |
| LRRC75A\|XM_021007003.1\|augustus_masked-186-processed-gene-52.0 |
| SERPINF2\|XM_021007050.1\|maker-186-snap-gene-61.44 |
| RPH3AL\|XM_021006972.1\|maker-186-snap-gene-64.24 |
| CERS2\|XM_021004679.1\|fragGeneModel-186-gene43-frag1 |
| HMGCS2\|XM_021003272.1\|maker-186-snap-gene-118.31 |
| maker-186-snap-gene-125.14 |
| MAB21L3\|XM_021002566.1\|maker-186-augustus-gene-132.4 |
| NRAS\|XM_021002322.1\|maker-186-augustus-gene-138.12 |

| |
|---|
| BCAS2|XM_021002294.1|maker-186-augustus-gene-139.8 |
| WNT2B|XM_021001840.1|maker-186-augustus-gene-146.14 |
| maker-186-augustus-gene-151.11 |
| KIAA1324|XM_021000574.1|maker-186-augustus-gene-160.35 |
| NTNG1|XM_021000241.1|augustus_masked-186-processed-gene-167.3 |
| OLFM3|XM_021000128.1|maker-186-augustus-gene-192.4 |
| VCAM1|XM_020999942.1|maker-186-augustus-gene-194.11 |
| LOC110217150|XM_020999329.1|maker-187-snap-gene-0.9 |
| LOC110200389|XM_020975685.1|maker-187-augustus-gene-79.3 |
| GLIS3|XM_021006269.1|fragGeneModel-187-gene20-frag1 |
| INSC|XM_021005437.1|maker-188-snap-gene-8.18 |
| CALCB|XM_021005540.1|augustus_masked-188-processed-gene-8.2 |
| COPB1|XM_021005410.1|maker-188-snap-gene-9.9 |
| NRIP3|XM_021002028.1|maker-188-augustus-gene-14.19 |
| LYVE1|XM_021001992.1|maker-188-snap-gene-20.15 |
| E2F8|XM_021001889.1|maker-188-augustus-gene-32.14 |
| NAV2|XM_021001949.1|fragGeneModel-188-gene21-frag1 |
| GAS2|XM_021001994.1|maker-188-snap-gene-45.9 |
| ANO3|XM_020977793.1|maker-188-snap-gene-54.9 |
| BDNF|XM_020977976.1|maker-188-snap-gene-57.11 |
| RCN1|XM_020963937.1|maker-188-augustus-gene-73.15 |
| MYBPC3|XM_021009684.1|maker-188-augustus-gene-125.38 |
| FAM180B|XM_021009830.1|maker-188-augustus-gene-126.29 |
| C1QTNF4|XM_021009823.1|maker-188-snap-gene-126.43 |
| DLGAP5|XM_020991147.1|fragGeneModel-189-gene46-frag2 |
| SEC24A|XM_020991221.1|maker-189-snap-gene-22.11 |
| PITX1|XM_020991397.1|maker-189-snap-gene-24.29 |
| POLE2|XM_021004787.1|maker-189-snap-gene-51.19 |
| TSHR|XM_020976319.1|maker-189-augustus-gene-137.2 |
| JARID2|XM_020986790.1|maker-190-augustus-gene-12.9 |
| ASIC1|XM_020966074.1|maker-191-augustus-gene-8.24 |
| C1QL4|XM_020966337.1|maker-191-augustus-gene-11.18 |
| LOC110194031|XM_020966140.1|fragGeneModel-191-gene4-frag1 |
| COL2A1|XM_020977216.1|fragGeneModel-191-gene13-frag1 |
| LRP8|XM_020987812.1|fragGeneModel-207-gene127-frag1 |
| DPH2|XM_020975471.1|fragGeneModel-207-gene35-frag1 |
| maker-208-snap-gene-15.15 |
| FHL2|XM_020987916.1|maker-214-snap-gene-10.19 |
| DYNC1I1|XM_020999971.1|fragGeneModel-218-gene5-frag1 |

| |
|---|
| SGCE\|XM_021000108.1\|maker-218-snap-gene-36.64 |
| maker-220-augustus-gene-31.2 |
| LOC110202722\|XM_020979006.1\|maker-220-snap-gene-66.25 |
| MYF6\|XM_020993512.1\|maker-221-snap-gene-10.19 |
| GMNN\|XM_020986850.1\|maker-224-snap-gene-37.26 |
| NTS\|XM_020993499.1\|maker-225-snap-gene-20.8 |
| DCN\|XM_021006426.1\|maker-225-snap-gene-40.22 |
| AMDHD1\|XM_021006465.1\|maker-225-snap-gene-55.9 |
| LOC110202709\|XM_020978976.1\|maker-225-augustus-gene-71.3 |
| PARPBP\|XM_020982353.1\|maker-225-augustus-gene-77.3 |
| ISL1\|XM_020993916.1\|maker-226-augustus-gene-13.8 |
| EMB\|XM_020996741.1\|maker-226-augustus-gene-18.2 |
| IGFBP3\|XM_020971545.1\|maker-227-snap-gene-9.15 |
| POC1A\|XM_020971566.1\|fragGeneModel-227-gene8-frag1 |
| ACY1\|XM_020999180.1\|maker-227-augustus-gene-23.27 |
| PCBP4\|XM_020998908.1\|maker-227-augustus-gene-23.24 |
| SLC38A3\|XM_020999235.1\|augustus_masked-227-processed-gene-31.0 |
| LOC110217133\|XM_020999308.1\|maker-227-augustus-gene-55.19 |
| REXO5\|XM_020965335.1\|maker-229-snap-gene-10.33 |
| C7\|XM_020996285.1\|maker-229-snap-gene-41.7 |
| DEPDC1B\|XM_021007291.1\|maker-229-augustus-gene-76.9 |
| LOC110199982\|XR_002322133.1\|augustus_masked-229-processed-gene-78.0 |
| CCNB1\|XM_020998473.1\|maker-229-augustus-gene-113.8 |
| LOC110201755\|XM_020977623.1\|maker-287-augustus-gene-1.23 |
| SLC39A4\|XM_020995164.1\|maker-350-augustus-gene-1.18 |
| GREM1\|XM_020995779.1\|augustus_masked-37-processed-gene-15.3 |
| maker-37-augustus-gene-28.11 |
| DUT\|XM_020976625.1\|fragGeneModel-37-gene41-frag1 |
| MAPK6\|XM_020980938.1\|fragGeneModel-37-gene50-frag1 |
| IGDCC3\|XM_020983201.1\|maker-37-snap-gene-130.37 |
| CILP\|XM_020983352.1\|maker-37-augustus-gene-130.30 |
| SNX1\|XM_020983684.1\|maker-37-snap-gene-134.25 |
| TICRR\|XM_020964953.1\|maker-37-snap-gene-220.32 |
| VPS33B\|XM_020965338.1\|maker-37-snap-gene-222.38 |
| FAH\|XM_020965802.1\|maker-37-augustus-gene-232.13 |
| MEX3B\|XM_020966003.1\|fragGeneModel-37-gene18-frag1 |
| EMX2\|XM_020988183.1\|maker-37-augustus-gene-315.2 |
| CPXM2\|XM_020964119.1\|maker-37-augustus-gene-346.25 |
| SLC16A12\|XM_020964159.1\|augustus_masked-37-processed-gene-346.10 |

| |
|---|
| CEP55\|XM_020964086.1\|maker-37-snap-gene-360.18 |
| RBP4\|XM_020964145.1\|maker-37-augustus-gene-360.16 |
| LGI1\|XM_020964188.1\|maker-37-augustus-gene-361.3 |
| HELLS\|XM_020964124.1\|snap_masked-37-processed-gene-363.3 |
| STAB2\|XM_020982366.1\|augustus_masked-379-processed-gene-4.3 |
| TDG\|XM_020982374.1\|maker-379-augustus-gene-4.28 |
| ALDH1L2\|XM_020982405.1\|maker-379-snap-gene-7.13 |
| CBLN2\|XM_020985630.1\|maker-38-augustus-gene-15.7 |
| IMPA2\|XM_020985559.1\|augustus_masked-38-processed-gene-67.0 |
| APCDD1\|XM_020985551.1\|maker-38-augustus-gene-73.9 |
| LAMA1\|XM_020975006.1\|maker-38-snap-gene-89.8 |
| ARHGAP28\|XM_020974993.1\|maker-38-snap-gene-89.9 |
| TYMS\|XM_020970658.1\|maker-38-snap-gene-114.10 |
| AQP4\|XM_020968730.1\|maker-38-snap-gene-140.9 |
| CHST9\|XM_020968675.1\|maker-38-augustus-gene-140.7 |
| CDH2\|XM_020992563.1\|fragGeneModel-38-gene52-frag1 |
| MCM4\|XM_020992585.1\|maker-38-augustus-gene-181.8 |
| PXDNL\|XM_020992584.1\|maker-38-snap-gene-199.14 |
| LYN\|XM_021005932.1\|maker-38-augustus-gene-215.4 |
| CPA6\|XM_021002092.1\|maker-38-snap-gene-262.14 |
| SULF1\|XM_021002089.1\|maker-38-augustus-gene-270.5 |
| MSC\|XM_020971030.1\|maker-38-augustus-gene-278.2 |
| GDAP1\|XM_020971036.1\|maker-38-snap-gene-287.2 |
| ZFHX4\|XM_020973940.1\|fragGeneModel-38-gene22-frag1 |
| STMN2\|XM_020976211.1\|maker-38-augustus-gene-312.9 |
| HEY1\|XM_020976213.1\|maker-38-augustus-gene-312.10 |
| CA2\|XM_020995738.1\|maker-38-augustus-gene-340.4 |
| SERPINB5\|XM_020995823.1\|maker-38-snap-gene-345.11 |
| EGFL6\|XM_021007515.1\|maker-419-snap-gene-16.5 |
| RBBP7\|XM_021007404.1\|maker-419-augustus-gene-31.11 |
| ADGRG2\|XM_021007455.1\|maker-419-augustus-gene-43.11 |
| LOC110214697\|XM_020995730.1\|fragGeneModel-42-gene8-frag1 |
| ATOH8\|XM_020969527.1\|maker-446-augustus-gene-12.32 |
| SFTPB\|XM_020969537.1\|maker-446-snap-gene-12.41 |
| KRT7\|XM_020966241.1\|maker-454-snap-gene-1.23 |
| LOC110194061\|XM_020966197.1\|snap_masked-454-processed-gene-4.18 |
| LOC110194060\|XM_020966194.1\|maker-454-augustus-gene-4.59 |
| HOXC5\|XM_020966246.1\|maker-454-snap-gene-8.46 |
| COPZ1\|XM_020966251.1\|maker-454-augustus-gene-10.34 |

| |
|---|
| GTSF1\|XM_020966248.1\|maker-454-augustus-gene-10.39 |
| LOC110192426\|XM_020963578.1\|snap_masked-460-processed-gene-0.0 |
| FAM105A\|XM_021008630.1\|maker-47-augustus-gene-41.7 |
| PARP11\|XM_021001653.1\|maker-470-snap-gene-6.19 |
| COPS7A\|XM_020994144.1\|maker-470-augustus-gene-25.100 |
| EBF3\|XM_020978347.1\|fragGeneModel-490-gene9-frag1 |
| CTBP2\|XM_020973635.1\|maker-490-augustus-gene-37.20 |
| maker-490-snap-gene-42.17 |
| CDK1\|XM_020998246.1\|fragGeneModel-490-gene19-frag1 |
| TMEM26\|XM_020998234.1\|maker-490-snap-gene-80.6 |
| IL11RA\|XM_020966933.1\|maker-520-augustus-gene-2.34 |
| CNN2\|XM_020969316.1\|maker-523-augustus-gene-6.30 |
| FAM32A\|XM_020969225.1\|maker-523-augustus-gene-13.7 |
| SEPT6\|XM_020975115.1\|maker-532-augustus-gene-2.28 |
| UBE2A\|XM_020975114.1\|maker-532-augustus-gene-2.31 |
| SPDL1\|XM_020992385.1\|maker-551-augustus-gene-38.9 |
| GABRB2\|XM_020992408.1\|fragGeneModel-551-gene25-frag1 |
| PTTG1\|XM_020992356.1\|maker-551-augustus-gene-75.10 |
| GALNT10\|XM_020978370.1\|maker-551-snap-gene-100.10 |
| G3BP1\|XM_020964239.1\|fragGeneModel-551-gene5-frag1 |
| STK32A\|XM_020964198.1\|maker-551-augustus-gene-121.7 |
| maker-551-snap-gene-124.16 |
| maker-558-snap-gene-14.54 |
| OSR1\|XM_020967455.1\|maker-559-snap-gene-8.20 |
| NRN1\|XM_020973828.1\|maker-582-augustus-gene-26.6 |
| ATP6V0D2\|XM_020970101.1\|maker-582-augustus-gene-50.13 |
| CUNH14orf180\|XM_020979473.1\|maker-64-snap-gene-18.6 |
| AURKB\|XM_020966832.1\|fragGeneModel-673-gene6-frag1 |
| BDKRB2\|XM_021007354.1\|maker-685-augustus-gene-1.8 |
| LOC110220791\|XM_021004983.1\|maker-685-augustus-gene-11.26 |
| CDC25C\|XM_021005047.1\|maker-685-snap-gene-12.44 |
| CHST2\|XM_020970967.1\|maker-688-augustus-gene-47.8 |
| LAMP3\|XM_020980833.1\|maker-688-augustus-gene-80.16 |
| FGF2\|XM_021008271.1\|maker-689-augustus-gene-11.25 |
| MAD2L1\|XM_021008357.1\|fragGeneModel-689-gene15-frag1 |
| RASL11B\|XM_020990071.1\|maker-76-snap-gene-35.12 |
| maker-76-snap-gene-112.2 |
| EDNRA\|XM_020994059.1\|maker-76-snap-gene-171.13 |
| GUCY1B3\|XM_020994058.1\|maker-76-augustus-gene-210.9 |

| |
|---|
| GRIA2|XM_020994112.1|maker-76-augustus-gene-218.10 |
| OIP5|XM_021009375.1|maker-77-augustus-gene-13.20 |
| COCH|XM_021004239.1|maker-77-snap-gene-45.9 |
| maker-77-snap-gene-50.17 |
| CCDC3|XM_020999731.1|fragGeneModel-79-gene87-frag1 |
| RINT1|XM_020984555.1|maker-84-snap-gene-2.17 |
| TES|XM_020979307.1|maker-84-snap-gene-46.6 |
| CEBPZ|XM_020964904.1|maker-87817-augustus-gene-0.6 |
| UAP1L1|XM_020981800.1|maker-90137-snap-gene-1.74 |
| HDGF|XM_020991496.1|fragGeneModel-90165-gene2-frag1 |
| SLC43A1|XM_020974509.1|maker-90195-augustus-gene-1.43 |
| NUP93|XM_020965598.1|fragGeneModel-90281-gene2-frag1 |
| CIAPIN1|XM_020965520.1|maker-90281-augustus-gene-3.30 |
| ILK|XM_020975852.1|fragGeneModel-90294-gene2-frag1 |
| HPX|XM_020975938.1|maker-90294-augustus-gene-3.44 |
| CHID1|XM_020979332.1|augustus_masked-90317-processed-gene-0.0 |
| maker-90330-augustus-gene-3.26 |
| BYSL|XM_020979741.1|maker-90352-augustus-gene-1.21 |
| MDFI|XM_020999646.1|augustus_masked-90352-processed-gene-3.0 |
| LOC110205079|XM_020981378.1|maker-90383-snap-gene-6.56 |
| DUSP9|XM_020977289.1|maker-90386-augustus-gene-3.20 |
| LOC110201392|XM_020976983.1|augustus_masked-90407-processed-gene-4.3 |
| DTL|XM_021007895.1|maker-90434-snap-gene-4.8 |
| NEK2|XM_021007868.1|maker-90434-augustus-gene-5.11 |
| LOC110195014|XM_020967665.1|maker-90444-snap-gene-7.61 |
| GCNA|XM_020976620.1|maker-90451-augustus-gene-4.32 |
| NVL|XM_020982180.1|augustus_masked-90466-processed-gene-6.2 |
| PCIF1|XM_020965726.1|maker-90503-snap-gene-8.93 |
| SIM2|XM_020996068.1|maker-90515-augustus-gene-3.19 |
| SRPX|XM_020979630.1|maker-90574-augustus-gene-20.18 |
| CTNNBL1|XM_020990760.1|maker-90591-snap-gene-4.8 |
| F8|XM_020975577.1|maker-90601-augustus-gene-9.25 |
| SLITRK1|XM_020976373.1|fragGeneModel-90602-gene2-frag1 |
| RBP1|XM_020986138.1|maker-90604-augustus-gene-7.10 |
| KPNA2|XM_021000781.1|fragGeneModel-90617-gene8-frag1 |
| CD7|XM_021000862.1|augustus_masked-90617-processed-gene-5.4 |
| ASPSCR1|XM_021000823.1|fragGeneModel-90617-gene11-frag1 |
| NPB|XM_021000892.1|maker-90617-augustus-gene-7.26 |
| LOC110218191|XM_021000758.1|fragGeneModel-90617-gene6-frag1 |

| |
|---|
| IGF2BP1\|XM_020997779.1\|maker-90630-augustus-gene-8.24 |
| EXO1\|XM_020982068.1\|maker-90633-snap-gene-7.12 |
| OPN3\|XM_020982058.1\|maker-90633-augustus-gene-7.7 |
| FH\|XM_020982053.1\|fragGeneModel-90633-gene4-frag1 |
| ZNF804A\|XM_020978671.1\|maker-90638-augustus-gene-17.19 |
| CCNA1\|XM_020967861.1\|maker-90639-augustus-gene-22.4 |
| RHOU\|XM_021000447.1\|maker-90650-augustus-gene-20.2 |
| CCSAP\|XM_021000449.1\|maker-90650-augustus-gene-22.5 |
| AGT\|XM_021000473.1\|maker-90650-snap-gene-28.26 |
| SEMA5B\|XM_020985956.1\|maker-90654-snap-gene-4.11 |
| HACD2\|XM_020985978.1\|maker-90654-augustus-gene-9.9 |
| UMPS\|XM_020985995.1\|maker-90654-augustus-gene-13.9 |
| SLC51A\|XM_020986039.1\|maker-90654-snap-gene-21.14 |
| LOC110208182\|XM_020986065.1\|maker-90654-snap-gene-23.29 |
| LOC110192040\|XM_020963072.1\|fragGeneModel-90667-gene2-frag1 |
| LOC110203353\|XM_020979830.1\|maker-90681-snap-gene-1.34 |
| IL13RA2\|XM_020963491.1\|maker-90681-augustus-gene-8.9 |
| CENPI\|XM_020963422.1\|maker-90681-augustus-gene-16.37 |
| SRPX2\|XM_020963444.1\|maker-90681-augustus-gene-17.29 |
| TNMD\|XM_020963440.1\|maker-90681-augustus-gene-18.16 |
| LOC110192334\|XM_020963434.1\|augustus_masked-90681-processed-gene-35.1 |
| TDRP\|XM_020973009.1\|maker-90686-augustus-gene-0.5 |
| ANKRD45\|XM_020970455.1\|maker-90692-snap-gene-28.22 |
| PRDX4\|XM_020989892.1\|augustus_masked-90696-processed-gene-2.1 |
| MAOA\|XM_020995885.1\|maker-90700-augustus-gene-26.5 |
| LOC110214856\|XM_020995943.1\|maker-90700-augustus-gene-26.6 |
| RCAN2\|XM_020993898.1\|maker-90701-snap-gene-16.20 |
| CUNH6orf226\|XM_020976859.1\|augustus_masked-90701-processed-gene-31.0 |
| CLCF1\|XM_020968326.1\|maker-90721-snap-gene-13.66 |
| CHKA\|XM_020968219.1\|fragGeneModel-90721-gene11-frag1 |
| CPT1A\|XM_020968228.1\|maker-90721-augustus-gene-17.21 |
| LOC110197178\|XM_020970861.1\|maker-90721-snap-gene-25.22 |
| HMBS\|XM_020984816.1\|augustus_masked-90723-processed-gene-1.3 |
| IL10RA\|XM_020984765.1\|maker-90723-augustus-gene-5.3 |
| LOC110207174\|XM_020984762.1\|maker-90723-augustus-gene-6.11 |
| KIF5C\|XM_020984747.1\|fragGeneModel-90723-gene8-frag1 |
| PCDH9\|XM_020992426.1\|maker-90730-augustus-gene-7.7 |
| MID1\|XM_021006245.1\|fragGeneModel-90731-gene13-frag1 |
| FAM46A\|XM_020969918.1\|maker-90734-augustus-gene-5.12 |

| |
|---|
| TTK\|XM_020975522.1\|maker-90734-snap-gene-13.6 |
| COL12A1\|XM_021000133.1\|maker-90734-augustus-gene-40.4 |
| maker-90734-snap-gene-41.10 |
| AK6\|XM_021007555.1\|fragGeneModel-90734-gene11-frag1 |
| maker-90740-snap-gene-4.28 |
| TLL2\|XM_020997114.1\|maker-90740-augustus-gene-4.17 |
| CRTAC1\|XM_020996911.1\|maker-90740-snap-gene-9.15 |
| DRGX\|XM_020996774.1\|maker-90740-augustus-gene-21.9 |
| GDF10\|XM_020997071.1\|maker-90740-augustus-gene-29.8 |
| MMD\|XM_020991935.1\|maker-90750-augustus-gene-14.6 |
| CA10\|XM_020972736.1\|maker-90750-augustus-gene-32.19 |
| CACNA1G\|XM_020972560.1\|maker-90750-snap-gene-35.52 |
| CNTNAP1\|XM_020972594.1\|snap_masked-90750-processed-gene-39.35 |
| PSMC3IP\|XM_020972659.1\|maker-90750-augustus-gene-40.30 |
| P3H4\|XM_020972782.1\|maker-90750-augustus-gene-43.24 |
| CFAP221\|XM_020963130.1\|maker-90752-snap-gene-0.8 |
| TMEM37\|XM_020963114.1\|maker-90752-augustus-gene-1.14 |
| MARCO\|XM_020972928.1\|maker-90752-snap-gene-4.17 |
| MCM6\|XM_021009486.1\|augustus_masked-90752-processed-gene-39.2 |
| LOC110206394\|XM_020983723.1\|augustus_masked-90760-processed-gene-7.1 |
| CTPS1\|XM_020983449.1\|maker-90760-augustus-gene-35.14 |
| P3H1\|XM_020983417.1\|maker-90760-augustus-gene-42.47 |
| HOXD10\|XM_020991022.1\|maker-90763-snap-gene-14.26 |
| NOP58\|XM_020972824.1\|maker-90769-snap-gene-61.25 |
| SLC1A4\|XM_020985435.1\|maker-90775-snap-gene-29.20 |
| LOC110207694\|XM_020985443.1\|maker-90775-snap-gene-33.4 |
| OTX1\|XM_020985480.1\|maker-90775-augustus-gene-38.4 |
| ADSS\|XM_020982096.1\|maker-90778-snap-gene-17.13 |
| PMPCA\|XM_020981892.1\|maker-90779-snap-gene-1.35 |
| CACFD1\|XM_020981989.1\|maker-90779-augustus-gene-15.26 |
| maker-90779-augustus-gene-18.15 |
| CFAP77\|XM_020973544.1\|maker-90779-snap-gene-19.19 |
| ASB6\|XM_020973522.1\|maker-90779-augustus-gene-31.8 |
| UPK1B\|XM_020985854.1\|maker-90792-snap-gene-57.7 |
| maker-90804-augustus-gene-2.10 |
| maker-90804-snap-gene-24.34 |
| LOC110207269\|XM_020984898.1\|fragGeneModel-90804-gene13-frag1 |
| HSPD1\|XM_020993611.1\|fragGeneModel-90804-gene16-frag1 |
| SLC39A10\|XM_020993521.1\|maker-90804-augustus-gene-43.3 |

| |
|---|
| SMOC2\|XM_020972071.1\|fragGeneModel-90808-gene3-frag1 |
| maker-90819-snap-gene-0.3 |
| GCLC\|XM_020993856.1\|maker-90822-augustus-gene-19.7 |
| SERPINH1\|XM_020989540.1\|maker-90823-snap-gene-16.15 |
| LOC110210562\|XM_020989538.1\|fragGeneModel-90823-gene14-frag2 |
| WNT11\|XM_020989533.1\|maker-90823-augustus-gene-20.15 |
| SCAMP5\|XM_020994527.1\|maker-90825-augustus-gene-12.9 |
| LOC110213925\|XM_020994497.1\|maker-90825-snap-gene-14.33 |
| ADPGK\|XM_020994454.1\|maker-90825-augustus-gene-24.12 |
| PAOX\|XM_020994672.1\|maker-90825-snap-gene-34.32 |
| ERLIN1\|XM_020994632.1\|maker-90825-augustus-gene-38.14 |
| CRABP1\|XM_020976023.1\|augustus_masked-90825-processed-gene-57.4 |
| RCN2\|XM_020976013.1\|maker-90825-augustus-gene-61.11 |
| LOC110200089\|XM_020975208.1\|maker-90825-snap-gene-70.8 |
| COL13A1\|XM_020975183.1\|maker-90825-augustus-gene-70.7 |
| DDX21\|XM_020975174.1\|maker-90825-augustus-gene-73.18 |
| HERC4\|XM_020998165.1\|maker-90825-augustus-gene-75.33 |
| FANCL\|XM_020996216.1\|maker-90842-augustus-gene-84.13 |
| POLD2\|XM_020984077.1\|maker-90845-augustus-gene-5.15 |
| YKT6\|XM_020984073.1\|maker-90845-augustus-gene-5.14 |
| SCARA5\|XM_020983996.1\|fragGeneModel-90845-gene11-frag1 |
| CGREF1\|XM_020983863.1\|maker-90845-augustus-gene-26.39 |
| CAD\|XM_020983838.1\|augustus_masked-90845-processed-gene-26.4 |
| LTBP1\|XM_020972346.1\|maker-90845-snap-gene-54.11 |
| KCNF1\|XM_020992711.1\|maker-90845-augustus-gene-87.4 |
| SULF2\|XM_020984167.1\|maker-90847-snap-gene-5.13 |
| CSE1L\|XM_020984170.1\|maker-90847-snap-gene-10.19 |
| BCAS4\|XM_020984275.1\|maker-90847-augustus-gene-16.11 |
| KCNG1\|XM_020984217.1\|maker-90847-augustus-gene-17.7 |
| AURKA\|XM_020981412.1\|maker-90847-snap-gene-38.18 |
| CUNH20orf85\|XM_020981357.1\|maker-90847-snap-gene-47.11 |
| NPEPL1\|XM_020981468.1\|maker-90847-augustus-gene-49.9 |
| SLCO4A1\|XM_020981543.1\|maker-90847-augustus-gene-74.21 |
| COL9A3\|XM_020981548.1\|maker-90847-augustus-gene-74.25 |
| SLC17A9\|XM_020981555.1\|maker-90847-augustus-gene-75.8 |
| CUNH2orf15\|XM_020992841.1\|maker-90849-snap-gene-80.11 |
| LOC110194436\|XM_020966796.1\|maker-90865-augustus-gene-6.43 |
| ATAT1\|XM_020974235.1\|snap_masked-90865-processed-gene-12.19 |
| ATP6V1G2\|XM_020974067.1\|maker-90865-snap-gene-14.47 |

| |
|---|
| LY6G6F\|XM_020974321.1\|maker-90865-snap-gene-15.42 |
| PBX2\|XM_020974141.1\|maker-90865-snap-gene-17.36 |
| HMGA1\|XM_020999502.1\|maker-90865-augustus-gene-27.5 |
| SCUBE3\|XM_020999564.1\|fragGeneModel-90865-gene44-frag1 |
| FANCE\|XM_020999584.1\|maker-90865-augustus-gene-33.8 |
| FKBP5\|XM_020999591.1\|maker-90865-augustus-gene-34.10 |
| ARMC12\|XM_020999580.1\|maker-90865-snap-gene-34.13 |
| GLP1R\|XM_020999358.1\|maker-90865-snap-gene-51.14 |
| LOC110194982\|XM_020967611.1\|fragGeneModel-90879-gene3-frag1 |
| C5\|XM_020990573.1\|maker-90879-augustus-gene-12.9 |
| LOC110196560\|XM_020969869.1\|maker-90881-augustus-gene-35.6 |
| VGLL1\|XM_020971772.1\|maker-90881-augustus-gene-41.14 |
| LOC110215779\|XM_020997567.1\|maker-90887-snap-gene-3.45 |
| TOP2A\|XM_020997649.1\|maker-90887-augustus-gene-4.20 |
| PNMT\|XM_020997632.1\|maker-90887-augustus-gene-7.20 |
| EXOSC8\|XM_020967909.1\|fragGeneModel-90895-gene4-frag1 |
| PTPRC\|XM_021003049.1\|maker-90895-snap-gene-79.13 |
| FMOD\|XM_021003235.1\|maker-90895-augustus-gene-96.19 |
| PRELP\|XM_021002971.1\|maker-90895-augustus-gene-97.8 |
| ETNK2\|XM_021003059.1\|maker-90895-augustus-gene-100.23 |
| PPP1R15B\|XM_021003117.1\|fragGeneModel-90895-gene32-frag1 |
| EIF2D\|XM_021002940.1\|maker-90895-augustus-gene-110.13 |
| C4BPA\|XM_021003243.1\|maker-90895-snap-gene-114.30 |
| CBS\|XM_020973887.1\|fragGeneModel-90897-gene7-frag1 |
| CRYAA\|XM_020973829.1\|maker-90897-snap-gene-3.16 |
| MRPS6\|XM_020995896.1\|maker-90897-snap-gene-30.9 |
| LOC110214881\|XM_020995983.1\|maker-90897-snap-gene-38.25 |
| NCAM2\|XM_020996054.1\|maker-90897-augustus-gene-48.7 |
| PWP2\|XM_020995940.1\|fragGeneModel-90897-gene24-frag1 |
| IGFBP5\|XM_021002757.1\|maker-90903-augustus-gene-7.6 |
| NDUFS1\|XM_020972819.1\|maker-90903-augustus-gene-49.15 |
| WDR12\|XM_020972817.1\|maker-90903-augustus-gene-62.20 |
| ISM1\|XM_020986149.1\|maker-90919-snap-gene-17.3 |
| LOC110199975\|XM_020975025.1\|fragGeneModel-90919-gene10-frag1 |
| PAX1\|XM_020986943.1\|fragGeneModel-90919-gene24-frag1 |
| FBLN7\|XM_020987533.1\|maker-90919-augustus-gene-71.4 |
| ITPRIPL1\|XM_020988518.1\|maker-90919-snap-gene-84.19 |
| SFRP1\|XM_020969593.1\|maker-90919-augustus-gene-126.4 |
| IDO1\|XM_020979690.1\|maker-90919-snap-gene-132.11 |

| |
|---|
| ADGRA2\|XM_020985273.1\|maker-90919-augustus-gene-138.18 |
| CBX2\|XM_021000929.1\|maker-90925-augustus-gene-10.9 |
| LOC110215852\|XM_020997666.1\|snap_masked-90925-processed-gene-16.8 |
| MIF4GD\|XM_021002449.1\|snap_masked-90925-processed-gene-42.11 |
| GRIN2C\|XM_021002582.1\|maker-90925-augustus-gene-45.20 |
| SDK2\|XM_021002493.1\|maker-90925-augustus-gene-52.13 |
| FAM20A\|XM_021000870.1\|maker-90925-augustus-gene-76.15 |
| CENPW\|XM_020987176.1\|maker-90977-augustus-gene-73.6 |
| FABP7\|XM_020987174.1\|maker-90977-snap-gene-88.7 |
| FYN\|XM_021008091.1\|fragGeneModel-90977-gene57-frag1 |
| WASF1\|XM_021008066.1\|maker-90977-snap-gene-142.3 |
| LIN28B\|XM_020998334.1\|fragGeneModel-90977-gene46-frag1 |
| SIM1\|XM_020998343.1\|maker-90977-augustus-gene-186.31 |
| EPHA7\|XM_020963076.1\|fragGeneModel-90977-gene1-frag1 |
| LOC110216801\|XM_020998793.1\|fragGeneModel-90994-gene17-frag1 |
| ZNF346\|XM_020998786.1\|maker-90994-augustus-gene-4.7 |
| FGFR4\|XM_020998780.1\|maker-90994-snap-gene-5.24 |
| SAMHD1\|XM_020998817.1\|maker-90994-snap-gene-28.26 |
| PTPRT\|XM_020998692.1\|fragGeneModel-90994-gene15-frag1 |
| S100A9\|XM_020992026.1\|maker-91023-augustus-gene-21.45 |
| S100A8\|XM_020991419.1\|maker-91023-augustus-gene-21.49 |
| FLAD1\|XM_020991628.1\|maker-91023-augustus-gene-27.37 |
| LOC110198751\|XM_020973220.1\|augustus_masked-91041-processed-gene-47.2 |
| MFAP2\|XM_020972750.1\|maker-91041-augustus-gene-58.35 |
| SZRD1\|XM_020972915.1\|maker-91041-augustus-gene-59.7 |
| FBLIM1\|XM_020973192.1\|maker-91041-augustus-gene-61.21 |
| SRM\|XM_020974014.1\|fragGeneModel-91041-gene12-frag1 |
| DFFA\|XM_020974117.1\|maker-91041-augustus-gene-80.5 |
| LOC110198087\|XM_020972225.1\|maker-91041-augustus-gene-120.19 |
| LOC110198097\|XM_020972244.1\|maker-91041-augustus-gene-120.16 |
| DLL3\|XM_020972453.1\|maker-91041-snap-gene-137.52 |
| SARS2\|XM_020978281.1\|maker-91041-augustus-gene-138.64 |
| PPP1R14A\|XM_020978782.1\|maker-91041-augustus-gene-140.28 |
| CADM1\|XM_020989713.1\|fragGeneModel-91133-gene32-frag1 |
| NCAM1\|XM_020989749.1\|fragGeneModel-91133-gene38-frag1 |
| PPP2R1B\|XM_020989791.1\|maker-91133-augustus-gene-27.14 |
| GRIA4\|XM_020996161.1\|fragGeneModel-91133-gene61-frag1 |
| LOC110215005\|XM_020996175.1\|augustus_masked-91133-processed-gene-57.0 |
| KIRREL3\|XM_021000632.1\|fragGeneModel-91133-gene75-frag1 |

| |
|---|
| FLI1\|XM_020963778.1\|maker-91133-augustus-gene-143.6 |
| SKA3\|XM_020968786.1\|maker-91133-augustus-gene-181.6 |
| FGF9\|XM_020968777.1\|maker-91133-augustus-gene-184.3 |
| SHISA2\|XM_020997241.1\|maker-91133-augustus-gene-196.6 |
| ALOX5AP\|XM_020997229.1\|maker-91133-augustus-gene-218.2 |
| CACNB4\|XM_020970344.1\|fragGeneModel-91133-gene12-frag1 |
| GPD2\|XM_021004861.1\|maker-91133-augustus-gene-273.5 |
| maker-98-augustus-gene-3.17 |
| COL22A1\|XM_020967066.1\|fragGeneModel-98-gene11-frag1 |
| LRRC6\|XM_020997907.1\|fragGeneModel-98-gene46-frag1 |
| NOV\|XM_020975547.1\|augustus_masked-98-processed-gene-120.0 |
| YWHAZ\|XM_020969456.1\|fragGeneModel-98-gene19-frag1 |
| CHEK2\|XM_020996586.1\|maker-99-augustus-gene-13.15 |
| CRYBB2\|XM_020996660.1\|maker-99-snap-gene-27.16 |
| TBX3\|XM_021007970.1\|maker-99-snap-gene-60.8 |

**Table S5.**
Summary of GO Biological Process terms enriched among PM8 genes.

See attached excel document

**Table S6.**
PM8 genes belonging to enriched Wnt-signaling GO term GO:0030111.

See attached excel document

**Table S7.**
Differentially expressed genes between the sugar glider patagium and dorsal skin prior to outgrowth.

See attached excel document

**Table S8.**
Differentially expressed genes between the sugar glider patagium and shoulder skin prior to outgrowth.

See attached excel document

**Table S9.**
Differentially expressed genes between outgrowing patagium and dorsal skin in the sugar glider.

See attached excel document

**Table S10.**
Differentially-expressed genes between outgrowing plagiopatagium and dorsal skin in Seba's short-tailed bat.

See attached excel document

**Table S11.**
Summary of GO Biological Process terms enriched among genes upregulated in the sugar glider patagium during its early outgrowth.

See attached excel document

**Table S12.**
Sugar glider patagium-upregulated genes belonging to enriched limb development GO terms.

See attached excel document

**Table S13.**

Summary of GO Biological Process terms enriched among genes upregulated in the bat plagiopatagium.

See attached excel document

**Table S14.**

Gene Ontology term enrichment among genes upregulated during plagiopatagium outgrowth in the bat.

See attached excel document

**Table S15.**

Differentially expressed genes shared between sugar glider and bat lateral patagia during their outgrowth.

| Gene ID | Differential Expression |
|---|---|
| ATRNL1 | Upregulated |
| BARX2 | Upregulated |
| BCL2 | Upregulated |
| CCDC3 | Upregulated |
| CREB5 | Upregulated |
| EPHA3 | Upregulated |
| ETV5 | Upregulated |
| GABRA1 | Upregulated |
| GABRB2 | Upregulated |
| GATA6 | Upregulated |
| GFRA1 | Upregulated |
| GREM1 | Upregulated |
| HAND2 | Upregulated |
| HAS2 | Upregulated |
| HECTD2 | Upregulated |
| HGF | Upregulated |
| ITGA4 | Upregulated |
| ITGA8 | Upregulated |
| MAN1A1 | Upregulated |
| OSR1 | Upregulated |
| PDE3A | Upregulated |
| TBX1 | Upregulated |
| TBX3 | Upregulated |
| TBX5 | Upregulated |
| TMEM26 | Upregulated |
| WNT5A | Upregulated |
| CLDN11 | Downregulated |
| FABP7 | Downregulated |
| FMOD | Downregulated |
| TENM2 | Downregulated |

**Movie S1.**

**Cell proliferation in sugar glider pouch young.** Light-sheet microscopy of postnatal day 3 sugar glider pouch young injected with EdU and collected 2hrs later. Visualization of cell proliferation (white signal) was carried using a modified iDISCO and tissue clearing protocol.
